# Heritable diel energy reserves enhance diatom growth

**DOI:** 10.64898/2026.01.03.697458

**Authors:** Oliver Müller, Tommaso Redaelli, Dieter A. Baumgartner, Clara Martínez-Pérez, Francesco Carrara, Johannes M. Keegstra, Roman Stocker

## Abstract

Diatoms are important drivers of marine primary production and biogeochemistry^1–4^. Compared to other phytoplankton groups, diatoms divide more asynchronously (i.e., divisions occur in a manner that is less aligned with the diel cycle) under non-limiting conditions, resulting in divisions occurring during both day and night^5–16^. However, the mechanisms and rates of asynchronous division have remained elusive. Here, using microfluidics-based time-resolved cell tracking, we measure the growth dynamics of individual cells of the diatom *Thalassiosira pseudonana* and show that cells growing mostly during the dark phase achieve rapid generation times (8 hours), dividing as fast as cells growing fully in light. We found that this remarkable ability of rapid growth in the dark is a consequence of the light history of both the cell and its parent cell, as light history controls the amount of photosynthetic energy the cell has stored in the form of the polysaccharide chrysolaminarin when entering the night. Furthermore, a mathematical model of this mechanism yields an up to 14% increase in the daily asynchronous population growth rate compared to growth without diel energy reserves when nutrients are non-limiting. This results in an up to 17-fold predicted increase in cell abundance over a typical 10-day diatom bloom, the maximal growth advantage of heritable chrysolaminarin during exponential phases of growth. By directly demonstrating and quantifying the benefit of chrysolaminarin under different diel conditions, this work provides a mechanistic understanding of how heritable diel energy reserves contribute to the rapid growth of diatom populations in the ocean. Beyond the specific discovery of the role of heritable energy reserves, we anticipate that our experimental approach can be broadly utilized in phytoplankton research by providing a blueprint to study phytoplankton physiology and ecology at the single-cell level, revealing novel mechanisms normally obscured in bulk growth assays.

## Introduction

Photosynthesis drives the sequestration of atmospheric carbon into biomass, with marine phytoplankton accounting for half of global carbon fixation^1,3^, thus directly influencing atmospheric carbon levels and Earth’s climate. For all light-harvesting organisms, the diel cycle of light and dark phases generates a periodicity in the energy supply for photosynthesis. This imposes a fundamental constraint on the growth of photosynthetic organisms and requires them to coordinate cellular processes with the fluctuating diel energy supply. Phytoplankton can fundamentally adopt two distinct diel growth strategies: dividing synchronously or asynchronously (also described as less-synchronized or less-entrained) with the diel cycle^13,15,17^. In synchronized populations, the cell cycle is strongly entrained by the light/dark cycle, with all cells within a population divide during one or sometimes two defined time windows of the diel cycle and, importantly, the population division rate is zero outside of these windows^13^. Synchronized divisions represent a low-risk growth strategy as they align energy production, nutrient uptake and cell division with the diel cycle, minimizing the risk of division during unfavourable times of the day^15,18,19^. Yet, synchronized divisions also limit the population’s growth rate, a significant trade-off, especially in seasonal summers where days are long and energy supply is abundant. Conversely, asynchronous divisions are weaker entrained by the diel cycle and instead multiple divisions occur throughout the light/dark cycle, enabling rapid population growth. In asynchronous populations there is always a fraction of the population dividing at any given point in the diel cycle and the population division rate is always greater than zero^13^. However, asynchronous divisions impose constraints on the metabolic fluxes of cells, as growth and divisions must occur in both light and darkness. When nutrients and light are abundant, asynchronous divisions enable rapid population growth, contributing to dense blooms that impact oceanic primary production^20–22^ which are expected to be strongly impacted by climate change^23–25^. Despite this importance, the mechanisms of asynchronous phytoplankton growth in relation to the diel light/dark cycle remain poorly understood.

Diatoms are among the most productive phytoplankton^1–4,26^, often attaining the highest growth rates among all phytoplankton groups^20,21,27,28^, dividing asynchronously under non-limiting conditions^5,7,9,11,13–16^. This has established asynchrony as a widespread paradigm for diatom growth^8,10^ and has led researchers to devote effort in creating synchronisation methods^6^. There have been occasional observations of synchrony in diatom growth in natural environments, but these are exceptions and are typically due to nutrient limitation^11^. Many aspects of diatom physiology—such as photosynthesis and cell cycle progression—are tightly linked to the diel cycle through both direct light cues^29–31^ and circadian rhythms^32–35^, which regulate diel transcriptional activity and coordinate cellular processes relative to the diel cycle. However, despite this strong diel coupling of many physiological processes, and the fact that diatoms can be temporarily forced into synchronous divisions through nutrient or light starvation^6,29,30^, diatom cell division remains largely asynchronous under nutrient-replete conditions^5–16^, with divisions occurring during both day and night (for an in-depth discussion and overview of diatom growth dynamics in light/dark cycles, see Supplementary Text). While asynchrony is not exclusively limited to diatoms and can occur e.g. during the germination of cyanobacteria resting spores^36^, compared to other major phytoplankton groups^37–60^, diatoms divide more asynchronously^5–16^ (i.e., divisions occur in a manner that is less aligned with the diel cycle) under non-limiting conditions, resulting in divisions occurring during both day and night. This leads to substantial variability in the light conditions experienced by individual cells within a population: some complete a cell cycle entirely in the light, whereas others do so almost entirely in darkness. Substantial progress has been made in elucidating the molecular mechanisms of light-dependent cell cycle checkpoints^29,30,61^ and circadian rhythms^33^, which ultimately commit cells to division within the diel cycle. Yet, how cells committed for division continue their growth during dark phases has remained mostly unknown. Pioneering studies have revealed diel patterns in energy storage compounds such as polysaccharides^62,63^, suggesting their importance in fuelling diel growth behaviour, but the effect of such compounds on single-cell growth rates has remained unknown. Therefore, how fast diatom cells can grow and divide during darkness has never been quantified and the mechanisms through which diatoms fuel asynchronous divisions remain unclear, limiting our understanding of how asynchrony contributes to the rapid growth of diatom populations.

Here, we report how the generation time of individual cells within an asynchronously growing population of the model diatom *Thalassiosira pseudonana* depends on the timing relative to the light/dark cycle. Using a novel experimental platform to study phytoplankton growth that combines microfluidics, time-lapse microscopy, and automated image analysis, we systematically tracked the growth trajectories (defined as the increase in cell size over a division cycle), measured the generation times (defined as the time for a complete division cycle, starting and ending with separation of newly formed daughter cells), and determined the lineage of thousands of individual *T. pseudonana* cells, for different light/dark cycles. We discovered that cells born during light can maintain rapid growth during the ensuing darkness, dividing as fast as cells growing exclusively in light. Analysis of diel polysaccharide levels showed that the ability to maintain a high growth rate in darkness results from the utilization of energy stored by both the cell and its parent cell in the form of the polysaccharide chrysolaminarin. Because chrysolaminarin (which is stored within the vacuole) gets partitioned during mitosis and shared across generations, we refer to it as a heritable diel energy reserve. Using a new mathematical model of single-cell diatom growth dynamics, we predict the relative contribution of such heritable diel polysaccharide reserves to asynchronous growth rates of diatom populations in different diel light/dark cycles. These findings contribute to explain how diatoms achieve rapid growth through asynchronous divisions—a process central to their contribution to global primary production and carbon cycling, through which they ultimately influence Earth’s climate and contribute to sustaining marine ecosystems.

## Results

### Single-cell generation times under continuous illumination

To obtain high-throughput measurements of the growth rates of single diatom cells, we developed a microfluidics-based experimental platform that integrates time-lapse microscopy and image analysis. We used this platform to determine how diel light/dark cycles shape individual cell growth in the model diatom *T. pseudonana*. Cells from batch flask cultures in early exponential phase (Extended Data Fig. 1a) were loaded into a microfluidic device comprising a single long microchannel (800 µm width × 200 µm height × 32 cm length), then mounted on an inverted microscope (Fig. 1a). Different diel light/dark cycles (continuous illumination: 24 h light; ‘long day’: 14 h light / 10 h dark; ‘short day’: 6 h light / 18 h dark) were generated by a circular ring light on the microscope connected to a timer, providing the actinic light (100 μmol photons m^−2^ s^−1^ of photosynthetically active radiation between 400 and 700 nm) to drive photosynthetic energy production (Fig. 1a, Extended Data Fig. 1b).

**Fig. 1.**
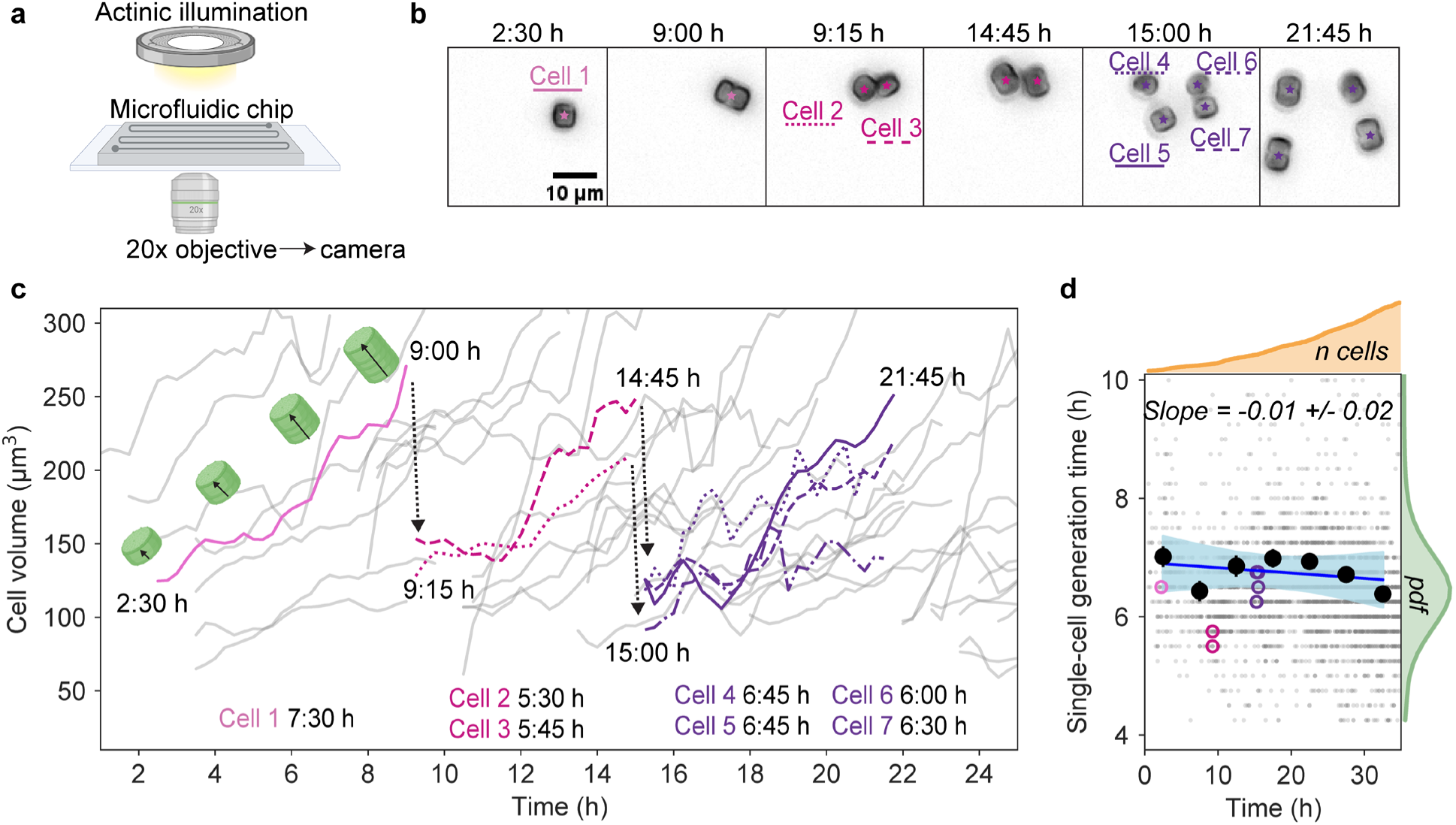
A microfluidics-based platform enables high-throughput tracking of single-cell diatom growth and division. **(a)** Schematic of the microfluidic setup used to quantify single-cell growth of the diatom *T. pseudonana*. **(b)** Representative time series of bright-field images showing a single *T. pseudonana* cell and its lineage across two generation within the microfluidic chip under continuous illumination. Cells were imaged every 15 min and only selected frames are shown (see Supplementary Video 1 for example of complete time series). Cell 1 grows between 2:30 (h:min) and 9:00, divides between 9:00 and 9:15 to form cells 2 and 3. Cells 2 and 3 grow between 9:15 and 14:45 and themselves divide between 14:45 and 15:00 to form cells 4–7, which grow between 15:00 and 21:45. Colors of the dots in each cell and associated cell number refer to the growth trajectories of each cell in panel (c). **(c)** Growth curves of single diatom cells all tracked simultaneously (grey), with cells 1–7 from panel b highlighted with their respective colors and their division events identified by dashed arrows. Solid arrows indicate cell expansion along the length axis of a diatom cell, resulting in increased cell volume (*i.e.*, growth).The text underneath the growth curves gives the single-cell generation time (time between birth of parent cell and division into two daughter cells) for each of cells 1–7. **(d)** Generation time of individual cells as a function of time in the experiment (grey dots), under continuous illumination. The generation time averaged over all cells in a 5 h window of time is shown by black circles, along with a fitted linear regression (blue line) and the 95% confidence interval of the slope (blue shading). Generation times of cells 1–7 (also given in panel (c)) are indicated in their respective colours (see panel (b)). The number of cells (orange curve) grows over time due to division events, up to *n* = 4682 cells. The green curve shows the probability density function of generation time. The values of time in each panel refer to the time into the experiment (illumination is continuous).

We imaged cells over the entire duration of the experiment (typically 52 – 72 h) using bright-field time-lapse microscopy (20x objective, 2048×2048 pixel sCMOS camera) with a temporal resolution of 15 minutes between images. During each experiment, cells underwent multiple rounds of growth and division (Fig. 1b). We analysed the resulting time series of images using a custom image analysis pipeline (Methods) to track the increase in volume of individual cells and detect division events. This allowed us to compute the generation time (time from birth to division) of each individual diatom cell, typically for 300-3000 cells per experiment (Fig. 1c).

First, we tracked the growth of individual cells under continuous illumination (Supplementary Video 1) to establish a baseline of single-cell generation times in the absence of a light/dark cycle. This yielded a distribution of generation times with a mean of *T_G_* = 6.6 h (standard deviation σ = 1.4 h) (Fig. 1d, green line, *n* = 4682 cells). Cells were born continuously throughout the experiment, highlighting the asynchronous nature of population growth under continuous illumination (Fig. 1d, orange line). The generation time remained constant during the experiment (Fig. 1d, blue line), indicating that our platform provides stable growth conditions over the experimental timeframe and that cells did not experience resource limitation.

### A longer dark phase increases the variability in generation time

Using the results for continuous illumination as a reference, we investigated whether diel light/dark cycles increased variability in generation times of single cells from an exponentially growing *T. pseudonana* culture. We subjected cells to one of two additional light regimes: long day (14 h light / 10 h dark, Supplementary Video 2) and short day (6 h light / 18 h dark, Supplementary Video 3). The light intensity was kept the same as in the continuous illumination (Fig. 2a, inset). In the following, we call the light period ‘day’, the dark period ‘night’, the transition from dark to light ‘sunrise’ and the transition from light to dark ‘sunset’. We tracked single-cell growth over three complete diel cycles and included cells born within the first two diel cycles in the subsequent analysis (Methods). We separately confirmed that the average growth rate in the microfluidic platform matched that of cells growing in conventional batch culture (Extended Data Fig. 2a).

**Fig. 2.**
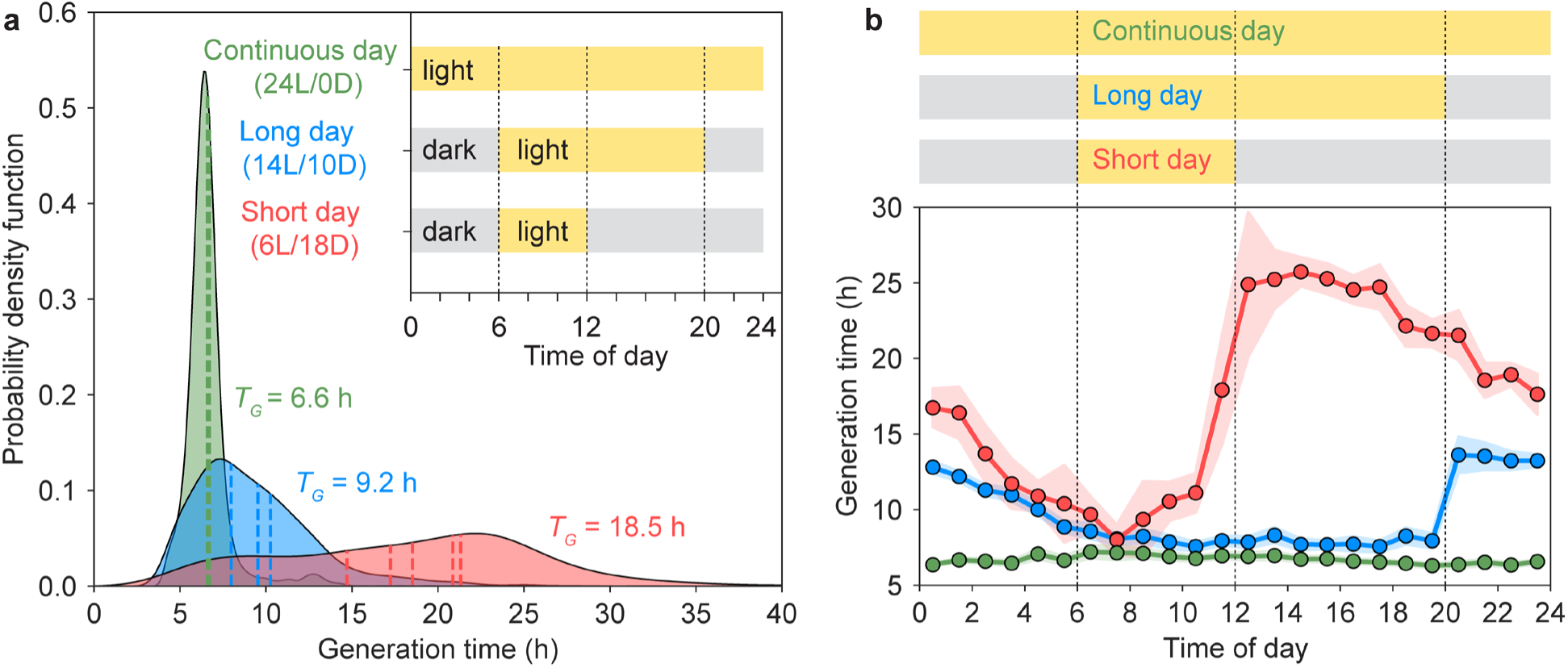
Diel light/dark cycle increase single-cell generation time variability in the marine diatom *T. pseudonana*. **(a)** Single-cell generation time distribution (probability density function, ‘pdf’) for cells grown under three different diel light (“L”) / dark (“D”) cycles (shown in inset): continuous day (green, 24 h light / 0 h dark, *n*_cells_ = 4682, *N* = 3 independent replicate experiment; data from Fig. 1d), long day (blue, 14 h light / 10 h dark, *n*_cells_ = 5455, *N* = 3 replicates) and short day (red, 6 h light / 18 h dark, *n*_cells_ = 1054, *N* = 5 replicates). Vertical dashed lines indicate the mean generation time of each replicate experiment. The values of *T_G_* refer to the mean generation time for each diel cycle. **(b)** Generation time of cells as a function of the time of day when the cell was born, for each diel cycle (see inset of panel (a) for the definition of time of day). The y-axis shows the generation time, i.e., the time it takes cells born at the time shown on the x-axis to undergo their next division. Circles show the mean of all cells within one-hour bins and shading represent the 95% confidence interval. Numbers of cells and replicates as in panel (a). The color of each curve corresponds to the color of the diel cycle represented above the panel and is the same as in panel (a).

Under continuous illumination, cells had a narrow distribution of generation times (mean *T_G_* = 6.6 h, standard deviation σ = 1.4 h, coefficient of variation CV = 0.21; Fig. 2a). For the long day, cells had a 40% higher mean generation time and a nearly three-fold wider distribution (*T_G_* = 9.2 h, σ = 3.8 h, CV = 0.38). For the short day, cells had a 200% higher mean generation time and a nearly five-fold wider distribution (*T_G_* = 18.5 h, σ = 7.6 h, CV = 0.40), compared to continuous illumination. These results show that the diel light/dark cycle impacts the variability of single-cell generation times in *T. pseudonana* and that longer nights lead to considerably slower growth rates. Despite the overall change in mean generation time, 23% of cells in the long day and 7% of cells in the short day still had a generation time equal to or smaller than the mean generation time under continuous illumination (6.6 h; Fig. 2a).

### Time of birth relative to the diel cycle is an important determinant of the generation time

Since diatom populations grow asynchronously^5–16^ (Extended Data Fig. 3), we hypothesized that the different spread in single-cell generation times within a population for different diel light/dark cycles stems from the different coupling of the timing of growth to the different diel cycles. To test this, we made use of the fact that our dataset allows us to correlate the timing of each cell’s growth trajectory to the time within the diel cycle. Specifically, we determined how the generation time of a cell depends on the timing of its birth within the diel cycle (Fig. 2b).

Under continuous illumination, the single-cell generation times (6.6 +/- 1.4 h) remained constant throughout the 24-hour period (Fig. 2b, green line), indicating no dependence of generation time on time of birth. In contrast, such a dependence existed in the long and short days. In the long day (Fig. 2b, blue line), cells born during the day grew rapidly, with generation times (7.9 +/- 3.1 h) close to those under continuous illumination. Generation times increased sharply after sunset, with cells born within 1 h after sunset having a 5.7 h longer generation time (13.6 +/- 3.1 h) than those born within 1 h before sunset (7.9 +/- 2.9 h). In the short day (Fig. 2b, red line), this effect was even more pronounced, with cells born within 1 h after sunset having a 7.0 h longer generation time (24.9 +/- 8.6 h) than those born within 1 h before sunset (17.9 +/- 10.5 h). These results provide direct evidence that a cell’s birth time relative to the light/dark cycle is an important determinant of its generation time. The presence of night introduced slow-growing, dark-born cells into the population and a longer night increased the generation time of these cells.

### Light history controls the growth of non-arrested cells during darkness

To further investigate the increase in the generation time across sunset in both long-day and short-day conditions (Fig. 2b), we examined the division behaviour of cells born before sunset (*-t_ss_*) and after sunset (*+t_ss_*) (Fig. 3b). Cells born before sunset (*-t_ss_*), and therefore in light, grew and divided during the subsequent dark phase without needing additional light, for both long-day and short-day conditions: we term these cells ‘non-arrested’. In contrast, cells born after sunset (*+t_ss_*), and thus in darkness, did not divide during the dark phase but waited for the next light period to divide: we term these cells ‘dark-arrested’ (Fig. 3a). This finding suggests that *T. pseudonana* has a light-dependent segment early in its cell cycle, requiring direct light exposure during that early time to eventually complete division.

**Fig. 3.**
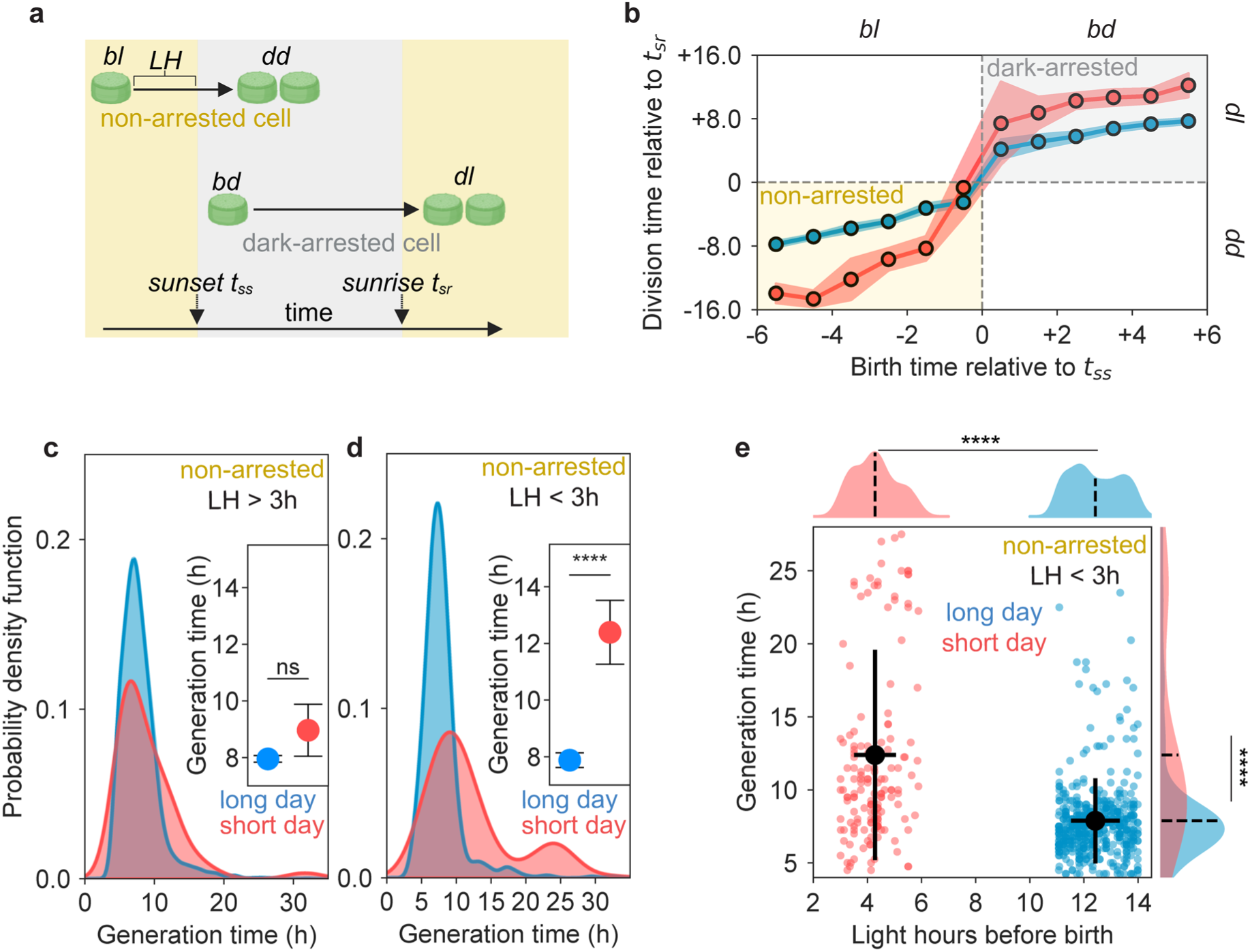
Light exposure of parent cells controls the growth of non-arrested cells during darkness. **(a)** Definition of non-arrested and dark-arrested cells. A ‘non-arrested’ cell is a cell born in light *(bl),* growing for a time (*LH)* in light and can divide during the ensuing dark phase *(dd)*. A ‘dark-arrested’ cell is a cell born in darkness *(bd)* and delaying division until at least the next light period (*dl*). Time of sunset is *t_ss_*, time of sunrise is *t_sr_*. **(b)** Cells born prior to sunset are non-arrested and divide during the subsequent dark phase, whereas cells born after sunset are dark-arrested and delay division until the next light phase. The time of division relative to sunrise (*t_sr_*) as a function of the time of birth relative to sunset (*t_ss_*), for the long day (blue) and short day (red). Cells born around sunset (*t_ss_*) can be classified into two groups: non-arrested *(bl + dd)* and dark-arrested *(bd + dl)*. The time of birth of cells was binned into 1 h bins. The mean (solid line) and 95% CI (shaded region) of the division time relative to *t_sr_* are shown. **(c, d)** Generation time distribution of non-arrested cells that received more than 3 hours of light prior to entering darkness (c, *LH* > 3) or less than 3 hours of light prior to entering darkness (d, *LH* < 3), for the long day (blue) and short day (red). (**c**) For cells with *LH* > 3, the diel cycle (long or short day) has no influence on the mean generation times (inset, mean ± 95% CI, comparison of means, *p* = 0.13, M.W.U test; *n*_long day_ = 2753 cells from three independent experiments; *n*_short day_ = 98 cells from five independent experiments). (**d**) For non-arrested cells with *LH* < 3, cells growing under short-day conditions have significantly longer generation times than under the long-day conditions (inset, mean ± 95% CI, *\*\*\*\* = p* < 0.0001, M.W.U test; *n*_long day_ = 496 from three independent experiments; *n*_short day_ = 157 cells from five independent experiments). **(e)** The light history experienced by progenitor cells (hours of light prior to cell birth) for non-arrested cells with *LH* < 3 (same cells as in panel (d)). Colored dots represent individual cells, for long-day(blue) or short-day (red) conditions, respectively. The mean (black solid circles in the main panel and black dashed lines in the outsets) and standard deviation (black solid lines) are shown. Compared to the short day, in the long day non-arrested cells receiving less than 3 h of light grow significantly faster (*p* < 0.0001, M.W.U test) and their progenitor cells received more light.

In another diatom, *Phaeodactylum tricornutum*, molecular studies showed that direct light exposure is required for the expression of a diatom-specific cyclin, a protein that controls the progression through the cell cycle, mediating the G1-to-S cell cycle transition^29,30^. While there is physiological evidence for this light dependency in the cell cycle of *T. pseudonana*^64^, whether the same molecular mechanism as in *P. tricornutum* occurs in *T. pseudonana* is not known.

However, given that many diatom-specific cyclins have also been identified in the genome of *T. pseudonana*^29^ and that we observed cells born during darkness to delay their division until the next day but those born just prior to darkness not to, we argue that a similar mechanism of light-dependent cell cycle checkpoints is likely to operate in *T. pseudonana* and to be responsible for the generation of ‘dark-arrested’ cells.

The generation time of dark-arrested cells between long-day and short-day conditions was significantly different (*p* < 0.0001, Mann–Whitney *U* (M.W.U) test, Extended Data Fig. 4a). This difference can be attributed primarily to the difference in the length of the dark phase experienced by arrested cells: the mean generation time difference of dark-arrested cells between long-day and short-day was 9.6 h, close to the 8-hour difference in the length of the night between the two conditions. Thus, cells born during the dark phase of the short-day condition experience a longer dark phase, delaying their division for longer.

Importantly, non-arrested cells also showed different generation times between the long-day and short-day conditions (*p* < 0.0001, M.W.U test, Extended Data Fig. 4b). Since non-arrested cells have passed the light-dependent cell cycle checkpoint, factors other than light-dependent cell cycle checkpoints must be at play to cause this difference.

The time of birth of non-arrested cells determines their light exposure, i.e. how many hours of light cells receive prior to division. We hypothesized that this light exposure, driving photosynthesis and energy production, is an important determinant of the generation time in non-arrested cells. To test this, we quantified the light hours (*LH*) experienced by a cell before darkness, under long-day or short-day conditions, and used this quantity to categorize cells into two groups: cells receiving more than 3 hours of light and those receiving less than 3 hours of light. Cells with *LH* < 3 h were born relatively shortly before sunset (Extended Data Fig. 4d), whereas those with *LH* > 3 h were born earlier in the light period (Extended Data Fig. 4c). Non-arrested cells that had received more than 3 hours of light had short generation times, with no significant difference between the short day and long day (*p* = 0.13, M.W.U test, Fig. 3c). In contrast, among cells that had received less than 3 hours of light, those from short-day conditions grew significantly slower than those from long-day conditions (*p* < 0.0001, M.W.U test, Fig. 3d), despite cells from both groups having received less than 3 h of light. This observation suggests that factors other than the light hours experienced by cells during the current generation contribute to the slower growth of cells receiving less than 3 hours of light under short-day conditions.

This observation led us to hypothesize that the light exposure of the progenitor cells might be important and underlie the growth difference in non-arrested cells with *LH* < 3 h between the long day and the short day (Fig. 3d). We therefore quantified the light hours experienced by the progenitor cells before a cell’s birth (for non-arrested cells with *LH* < 3 h; Fig. 3d). This analysis showed that progenitor cells in the long day had experienced significantly more light hours (*p* < 0.0001, M.W.U test) than their counterparts in the short day (Fig. 3e). Furthermore, cells with *LH* < 3 h had significantly shorter generation times (*p* < 0.0001, M.W.U test) in the long day (7.9 +/- 2.9 h) than in the short day (12.4 +/- 7.2 h). These results suggest that, in addition to the direct impact of a cell’s own light exposure, the light history of its progenitor cells influences the growth of non-arrested cells that received less than 3 h of light and thus grew mostly in darkness.

### The diel cycle controls the accumulation and consumption of chrysolaminarin

Non-arrested cells born near sunset (*LH* < 3 h), because they receive limited light, must perform most of the metabolic activity required for growth and division in darkness. In the absence of light, these cells rely on stored energy and carbon to sustain growth. We hypothesized that storage polysaccharides could support the growth of *T. pseudonana* cells during darkness, since population-averaged studies have shown that storage polysaccharides exhibit diel patterns of accumulation and consumption in both laboratory cultures and natural environments^62,63,65–67^.

To investigate the role of storage compounds in controlling generation times, we measured levels of chrysolaminarin, the primary polysaccharide storage compound in diatoms^68,69^, using aniline blue staining and quantification of the resulting fluorescence via flow cytometry^70^ (Fig. 4a).

**Fig. 4.**
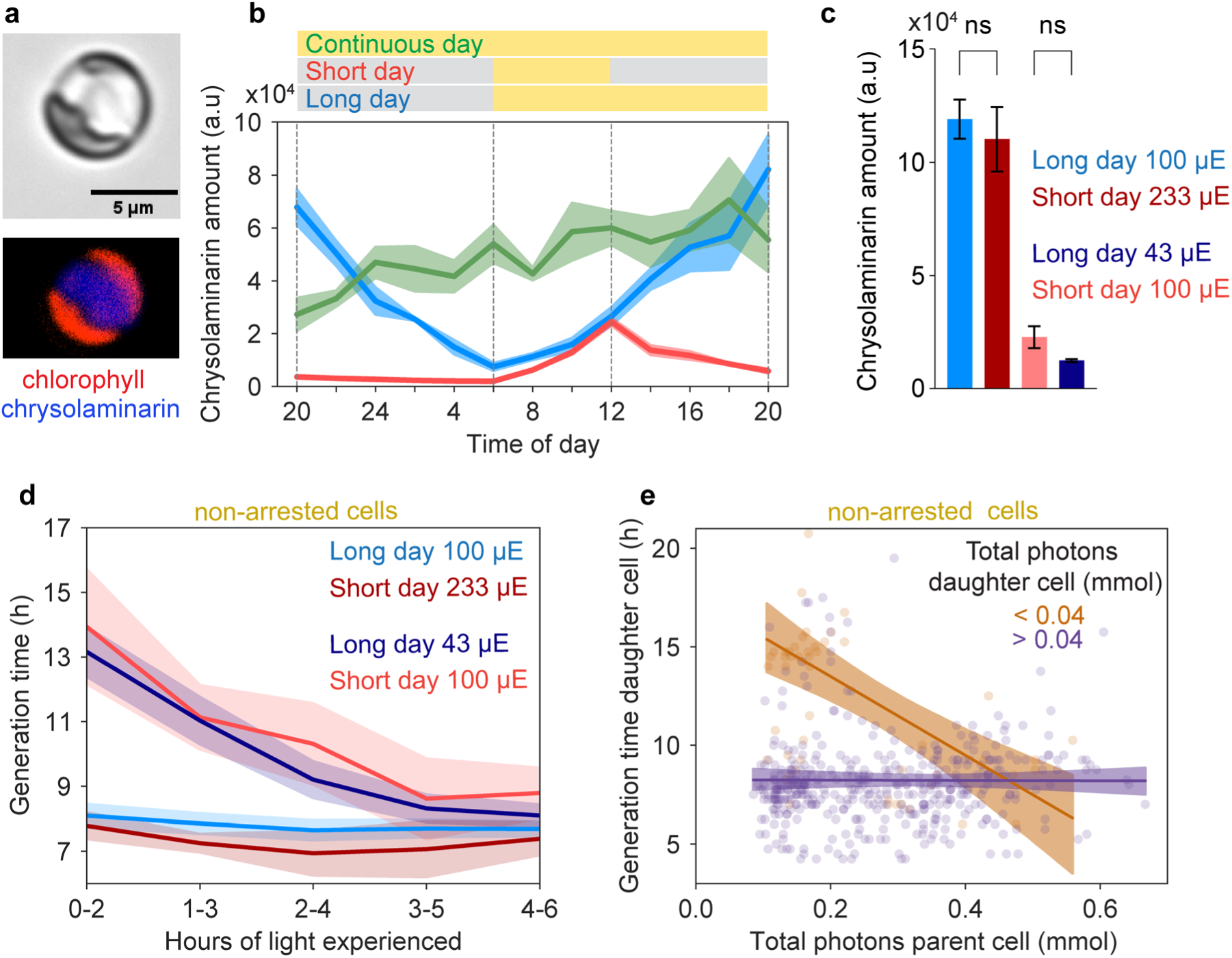
Light exposure controls chrysolaminarin accumulation and diel growth dynamics of non-arrested cells. **(a)** A *T. pseudonana* cell stained with aniline blue to visualize intracellular chrysolaminarin. Bright-field image (top) and fluorescence image (bottom), showing chlorophyll autofluorescence (red) and aniline-blue fluorescence (blue). **(b)** Diel pattern of chrysolaminarin storage (measured as chrysolaminarin fluorescence) under different diel cycles: continuous day (green), short day (red) and long day (blue). Chrysolaminarin levels were measured at 2 h intervals for a full 24 h diel cycle using flow cytometry (see Methods). Dashed lines indicate sunrise and sunset for the long-day and short-day conditions (see diel cycles above the image). Mean (solid line) and standard deviation (shaded area) are shown for three independent replicate experiments for each diel cycle. **(c)** Chrysolaminarin amount at sunset of cells growing in long-day or short-day diel cycles for different light intensities (43 µmol m^-2^ s^-1^, 100 µmol m^-2^ s^-1^ or 233 µmol m^-2^ s^-1^). All four diel conditions were compared to each other using one-way ANOVA and Tukey’s multiple comparison test, resulting in *p* < 0.0001 for all comparisons unless indicated as ns (non- significant). Means and standard deviations for four independent replicates per condition are shown. **(d)** Generation time of cells as a function of the number of hours of light they received, for cells growing in the long day at 100 µmol m^-2^ s^-1^ (blue, n_cells_ = 1464), in the short day at 100 µmol m^-2^ s^-1^ (red, n_cells_ = 232), in the long day at 43 µmol m^-2^ s^-1^ (dark blue, n_cells_ = 815) and in the short day at 233 µmol m^-2^ s^-1^ (dark red, n_cells_ = 470). Solid lines show a 2 h moving average and shading indicates the 95% CI of the mean. **(e)** The generation time of daughter cells as a function of the total photons received by their parent cell, for daughter cells which received less (orange, n_cells_ = 55) or more (violet, n_cells_ = 398) than 0.04 mmol photons prior to darkness. A Significant negative correlation exists for the former (Pearson correlation *r* =-0.49, *p* < 0.0001), but not the latter (*r* = -0.01, *p* = 0.93). Circles represent individual cells, solid lines the linear regressions and shading the 95% confidence interval.

Sampling cells from batch cultures at 2 h intervals for each of the three diel cycles showed a strong correlation between chrysolaminarin levels and the light/dark cycle (Fig. 4b). These data indicates that cells accumulated chrysolaminarin during the light period and degraded it during the dark period, so that levels were lowest at sunrise, increased throughout the day until sunset, at which point consumption began until the next sunrise (Fig. 4b). This observation suggests that chrysolaminarin serves as a diel energy reserve in *T. pseudonana*.

### Chrysolaminarin storage enables fast growth in darkness

To assess the impact of chrysolaminarin on the growth of *T. pseudonana*, we sought to further modulate chrysolaminarin accumulation and compare it to observed single-cell growth dynamics. Since both light duration and intensity can be used to modulate chrysolaminarin accumulation, we measured chrysolaminarin levels in cells exposed to either a long or short day at moderate light intensity (100 µmol m^-2^ s^-1^), a long day at reduced light intensity (43 µmol m^-2^ s^-1^) or a short day at increased light intensity (233 µmol m^-2^ s^-1^). The cumulative photons (light hours × light intensity) received until sunset in the long day at 43 µmol m^-2^ s^-1^ matched that in the short day at 100 µmol m^-2^ s^-1^ (2.16 × 10^-6^ mol photons m^−2^). In the long day at 100 µmol m^-2^ s^-1^, the cumulative photons received match the short day at 233 µmol m^-2^ s^-1^ and were 2.3-fold higher (5.03 × 10^-6^ mol photons m^−2^).

We found that the chrysolaminarin levels at the end of the light period depended on the cumulative photons received during the light period (Fig. 4c). Cells in the long day at 100 µmol m^-2^ s^-1^ and in the short day at 233 µmol m^-2^ s^-1^ had significantly more chrysolaminarin at sunset than those in the long day at 43 µmol m^-2^ s^-1^ or in the short day at 100 µmol m^-2^ s^-1^ (*p* < 0.0001, one-way ANOVA and Tukey’s multiple comparison test), whereas no difference was observed between short-day at 100 µmol m^-2^ s^-1^ and long-day at 43 µmol m^-2^ s^-1^ or between cells in the long day at 100 µmol m^-2^ s^-1^ and in the short day at 233 µmol m^-2^ s^-1^. These data show that cells in the long day at 100 µmol m^-2^ s^-1^ and in the short day at 233 µmol m^-2^ s^-1^ entered darkness with five times more chrysolaminarin than cells in the short day at 100 µmol m^-2^ s^-1^ and nine times more than cells in the long day at 43 µmol m^-2^ s^-1^.

We investigated whether these differences in chrysolaminarin levels at sunset translated to differences in generation times among the three diel conditions. Since we further hypothesized that cells born closer to sunset (and thus receiving less light) should depend more on stored energy, we analysed the generation time of non-arrested cells as a function of the light hours that non-arrested cells experienced before entering darkness (Fig. 4d). Cells which received less than 2 h of light showed the greatest difference in generation times between the long day at 100 µmol m^-2^ s^-1^ (8.1 +/- 3.1 h, *n* = 295), short day at 233 µmol m^-2^ s^-1^ (7.8 +/- 3.9 h, *n* = 221) and both the short day at 100 µmol m^-2^ s^-1^ (13.9 +/- 8.4 h, *n* = 86) and the long day at 43 µmol m^-2^ s^-1^ (13.2 +/- 5 h, *n* = 158). Furthermore, cells in the long day at 100 µmol m^-2^ s^-1^ and in the short day at 233 µmol m^-2^ s^-1^ had the same generation times (8.1 +/- 3.1 h, *n* = 295) independently of light hours received. In contrast, in the short day at 100 µmol m^-2^ s^-1^ generation times increased by 50% between cells receiving 4-6 h of light (8.8 +/- 3.3 h, *n* = 68) and those receiving 0-2 h of light (13.9 +/- 8.4 h, *n* = 86). A similar trend was observed for the long day at 43 µmol m^-2^ s^-1^, where generation times increased 1.6-fold (from 8.1 +/- 2.86 h, *n* = 279, to 13.2 +/- 4.95 h, *n* = 158). These results suggest that cells entering darkness with less chrysolaminarin grow slower during the ensuing night.

To assess the impact of the light exposure of a parent cell on the generation time of its daughter cell, we converted the light hours each parent cell experienced into total photons it received from birth to division (Methods) and correlated this quantity to the generation time of its daughter cell (Fig. 4e). For daughter cells that themselves received less than 0.04 mmol photons prior to darkness, we found that the generation time decreased with increasing amount of photons received by their parent cell (*r* = -0.49, *p* < 0.0001; Fig. 4e). In contrast, no dependence (*r* = - 0.01, *p* = 0.93) was observed for daughter cells that received more than 0.04 mmol photons prior to darkness. This result shows that cells receiving little light themselves can still grow fast if the total photons received by their parent cell is large.

Taken together, our results indicate that the generation time of a *T. pseudonana* cell depends on the cell’s chrysolaminarin quota, which is in turn determined by the cumulative light received by the cell and its parent cell. In the long-day 100 µmol m^-2^ s^-1^ and short-day at 233 µmol m^-2^ s^-1^ condition, cells born near sunset entered darkness with more chrysolaminarin, primarily accumulated and passed on by their parent cell, enabling rapid growth. In contrast, cells entering darkness with less chrysolaminarin because their parent cell had received less light (as in the short-day 100 µmol m^-2^ s^-1^ and long-day 43 µmol m^-2^ s^-1^ conditions) grew considerably slower. The increase in generation time of cells with low chrysolaminarin stores that are born closer to sunset reflects the increased dependency on stored energy (as opposed to energy production driven directly by sunlight) as cells have to perform more and more energy-requiring tasks during darkness. Our findings demonstrate that for cells receiving minimal light themselves, and thus performing most metabolic activities required for growth in darkness, chrysolaminarin inherited from parent cells is an important determinant of generation time. More generally, these results show that the population-averaged growth rate depends on the light history of both individual cells and their parent cells.

### Heritable chrysolaminarin storage provides a significant growth advantage to diatom populations

To quantify how heritable cellular chrysolaminarin storage affects overall diatom population growth, we developed an individual-based model of *T. pseudonana* growth based on the experimentally measured single-cell growth dynamics. The model, which we refer to as storage model, tracks cells as they progress through the cell cycle, culminating in division (*D*), and in parallel captures the accumulation and consumption of chrysolaminarin (Fig. 5a, Supplementary Methods). As in a previous model (‘Vaulot model’) of diel phytoplankton growth^71,72^, we represent the cell age through two variables: *n*, representing the cell age (in hours) in light-independent processes and *k*, representing the cell age (in hours) in light-dependent processes. The cell cycle of a new cell must complete a light-dependent segment (of length *k_1_* hours) and a light-independent segment (of length *n_1_* hours) (progression in *k_1_* and *n_1_* can occur simultaneously) before crossing a transition point (*T,* a light-dependent cell cycle checkpoint).

**Fig. 5.**
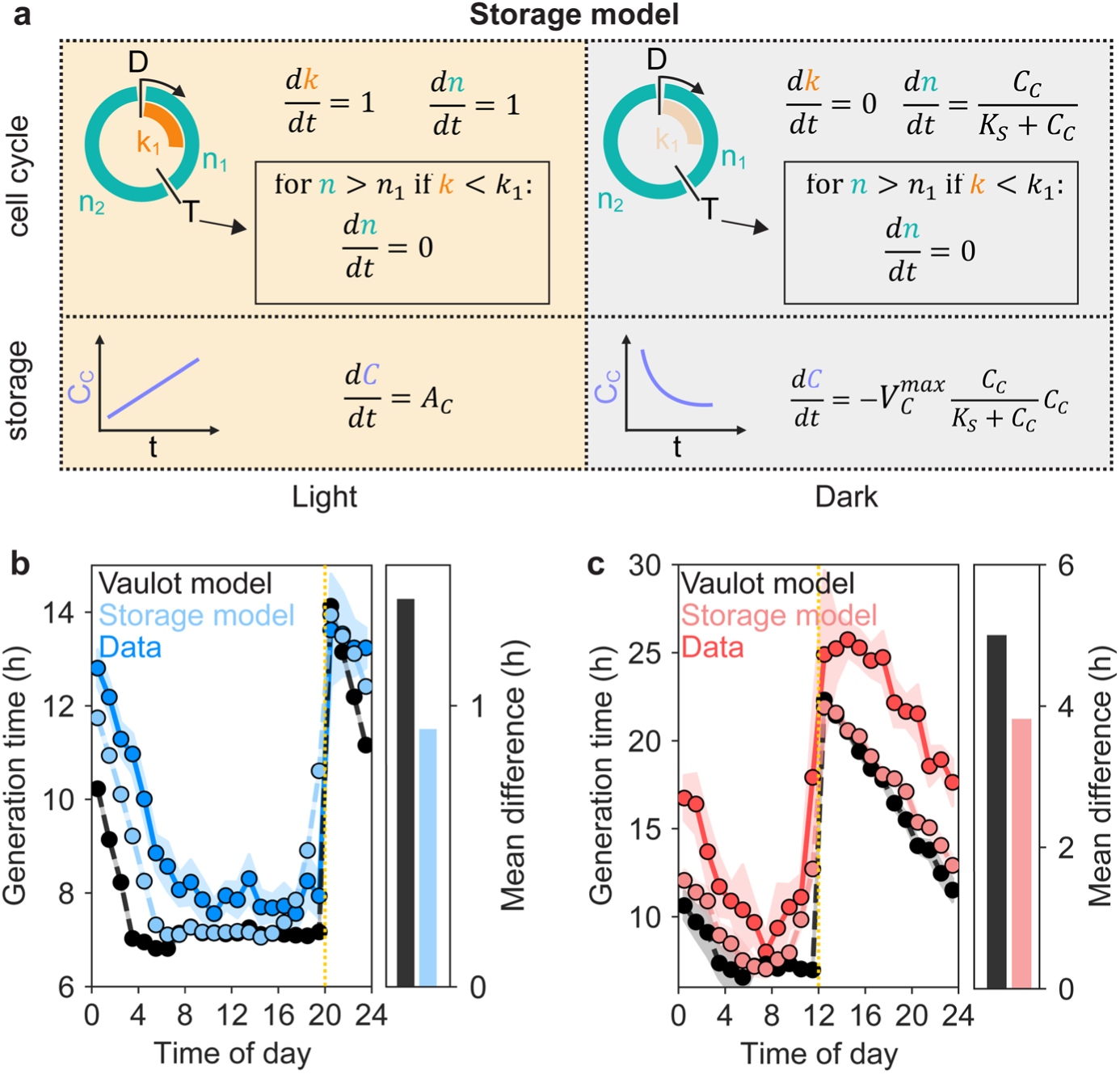
A mathematical model of diatom growth predicts the effect of heritable storage on diatom generation times. **(a)** Schematic of the cell cycle progression and chrysolaminarin dynamics in light and darkness used in our mathematical model (‘Storage model’) of diatom growth. To complete a full cell cycle, culminating in cell division (*D*), cells must complete one light-dependent (*k*_1_) and two light-independent (*n*_1_*, n*_2_) cell cycle segments. To cross the transition point (*T*) and start progressing in *n_2_* cells must first complete *k*_1_ and *n*_1_ (dashed arrow and black box). Cells can progress simultaneously in *k*_1_ and *n*_1_. After the transition point *T*, cells must complete *n*_2_ to divide. In light, cells progress at a constant rate (*dk/dt* = 1; orange, *dn/dt* = 1) in *k* and *n*. A cell’s chrysolaminarin level *C*_C_ in light increases with time at a rate *A*_C_. In darkness, cells cannot progress in *k* (*dk/dt* = 0; faint orange) but progress in *n* using stored chrysolaminarin, at a rate that depends on the amount of stored chrysolaminarin *C*_C_ via a Monod relationship with half-velocity constant *K*_S_. **(b, c)** Generation time of cells as a function of the time of day when the cell was born. Solid lines are the experimental data (from Fig. 2b) for the long day (solid line, dark blue in (b)) and short day (solid line, dark red in (c)). Colored dashed lines are the predictions of our model (Storage model) for the long day (dashed line, light blue in (b)) and short day (dashed line, light red in (c)). Black dashed lines (for both (b) and (c)) are the predictions of the classic model without storage (‘Vaulot model’). The vertical yellow dotted lines indicates sunset. Circles show the mean of 1-hour windows and shading indicates the 95% confidence interval. The bar plots on the right of the main panels give the mean difference between the experimental data and the predictions of the models with storage (blue in (b), red in (c)) and without storage (black, in both (b) and (c)), respectively.

After *T*, a cell must complete a second light-independent segment (of length *n_2_* hours) before it divides. Models based on this framework have been able to account for the light-dependent checkpoint-induced increase in generation time of cells born during darkness^71,72^ (arrest at *T* in darkness increases generation time). These models assume a constant rate of progression through light-independent and light-dependent processes (*n* and *k* always grow linearly in time) and therefore, cannot account for the dependency of non-arrested cells on storage, which we experimentally observed for cells born in the hours preceding sunset and thus growing mainly during darkness (see Fig. 5b,c).

Informed by our experimental single-cell growth trajectories, we therefore expanded on this framework by making the progression of *n* in darkness dependent on stored chrysolaminarin. We modelled cells as accumulating chrysolaminarin in light at a rate *A_C_* and consuming it to fuel the progression of the light-independent cell-cycle segments during darkness. The rate of chrysolaminarin consumption and chrysolaminarin-dependent progression through *n* follows a Monod relationship with a half velocity constant *K_S_,* assuming chrysolaminarin as the growth limiting substrate during darkness (Supplementary Methods). For a given light/dark cycle and set of cell parameters (*K_1_*, *N_1_*, *N_2_*, *A_C_*, *K_S_, T*, representing the expected length, in hours, of light dependent and light independent segments; Supplementary Methods), the model simulates single cells as they progress through *k* and *n* and ultimately divide into two daughter cells. During division, the chrysolaminarin store is split equally between the daughter cells. The model, which based on our experimentally observed increase in volume during a division cycle (Methods) assumes that cells double their biomass from birth to division, provides the birth times and generation times of all cells born during the simulation, from which we compute population-based metrics such as the mean growth rate.

To evaluate the model’s ability to capture the observed dependency of cells born before sunset on stored chrysolaminarin, we compared the generation times predicted by the model with our experimentally measured ones. The cell parameters used in the model were estimated based on our experimental data (Supplementary Methods, Extended Data Fig. 6). To determine the specific role of chrysolaminarin storage in the model predictions, we further added to the comparison the predictions of a previously developed model (Vaulot model)^71^, in which storage is not considered and cells progress at a constant rate through *n* and *k*.

Results show that our new model accounting for storage outperforms the earlier model neglecting storage in predicting the experimental data and in particular in capturing the dependence of non-arrested cells on stored chrysolaminarin, for both the long day (Fig. 5b) and the short day (Fig. 5c). For the long day, our model overestimates the generation time of cells born within 1 h of sunset, possibly due to underestimating chrysolaminarin stores available to cells close to sunset.

To determine the potential growth advantage afforded by chrysolaminarin storage in different environmental conditions, we ran the model with and without chrysolaminarin storage (*n* progression in darkness is chrysolaminarin-dependent vs. no progression of *n* in darkness as cells without chrysolaminarin cannot grow during the night) over a broad range of possible day/night cycles (from 6 to 22 h of darkness, with 0.5 hour increments) and light intensities (from 0 to 200 µmol m^-2^ s^-1^, with 20 µmol m^-2^ s^-1^ increments) (Fig. 6b, Supplementary Methods). For each of these environmental conditions we computed the daily mean of the population growth rate for both models (with and without chrysolaminarin storage). By then comparing the mean growth rate predicted by the two models, we determined the growth advantage afforded by chrysolaminarin storage. We found that the growth advantage strongly increases with longer nights For a moderate light intensity of 100 µmol m^-2^ s^-1^, the daily growth advantage increases from 8.8% for the shortest night (6 h) to 29.2% for the longest night (22 h). This three-fold increase is due to the increased benefit of storage compounds in long nights compared to short nights. When most growth in light-independent segments occurs in darkness, the relative benefit of faster growth via chrysolaminarin becomes larger. Furthermore, higher light intensities enhance this advantage by allowing cells and their parents to accumulate more chrysolaminarin ahead of darkness: The growth advantage in a long day with 14 h light / 10 h dark diel cycle increases from 8.5% to 14.2% as the light intensity increases from 10 to 200 µmol m^-2^ s^-1^.

**Fig. 6.**
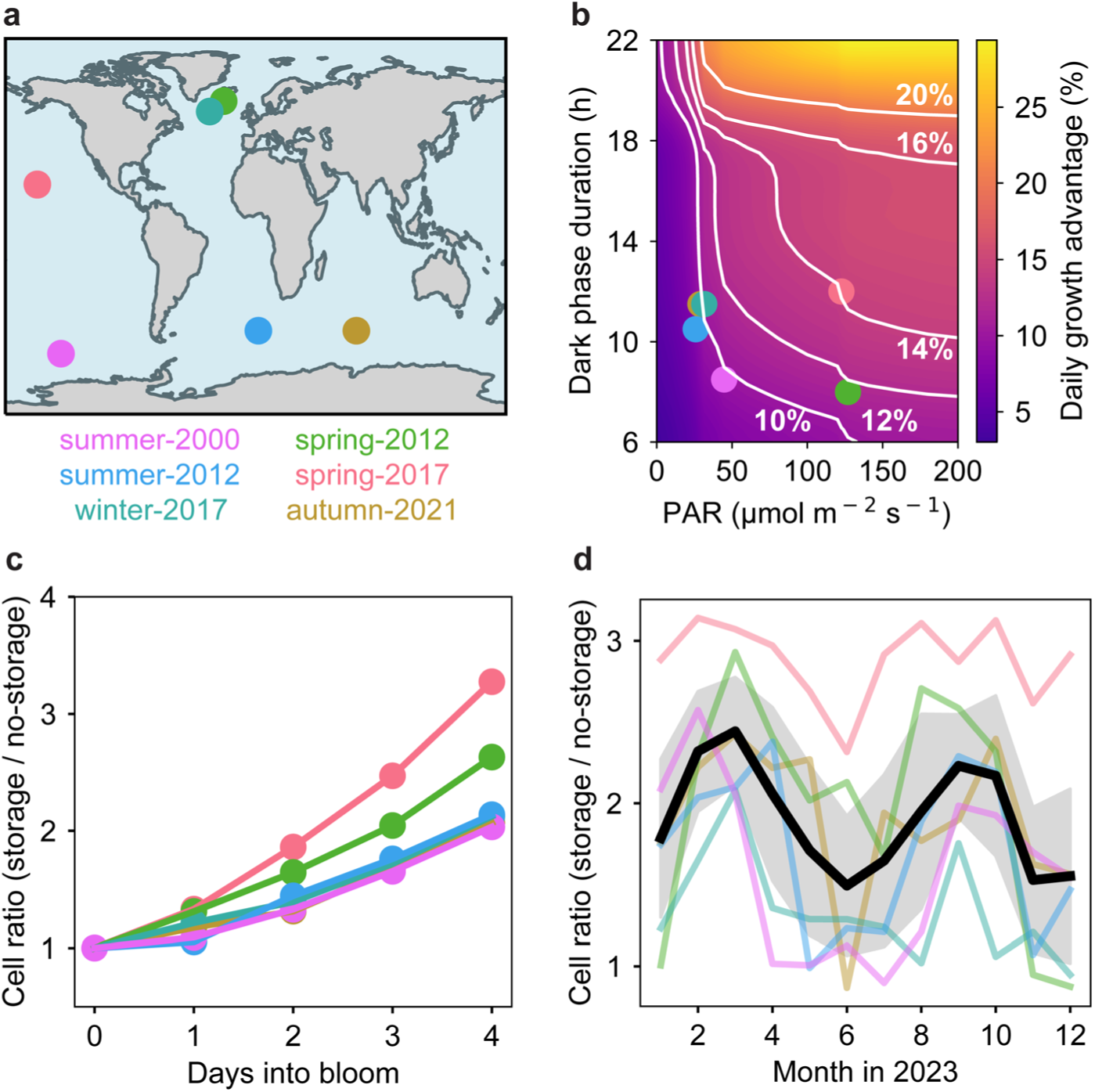
Heritable diel chrysolaminarin storage provides a considerable growth advantage to diatom populations. **(a)** Locations and times of six diatom blooms reported in the literature^73–78^, used to predict the effect of heritable energy storage through our model. **(b)** Daily growth advantage (colorbar) of a diatom population that uses chrysolaminarin storage, relative to one that does not, as predicted by the model, as a function of the dark phase duration and the light intensity (photosynthetically active radiation (PAR)). The white contour lines denote growth advantages of 10%, 12%, 14%, 16% and 20% per day. Environmental conditions for six diatoms blooms reported in the literature (see panel (a)) are indicated as colored circles. **(c)** Four-day time series of the size of a diatom population (number of cells) that uses chrysolaminarin storage, relative to one that does not, for the environmental conditions associated with the six blooms in panel (a), as predicted by our model. **(d)** Same, for environmental conditions associated with the six blooms in panel (a) for each month of the year 2023. For each month, the cell ratio is computed from our model at the end of a 4-day bloom. The black line represents the mean of the cell ratios for the six blooms and shading indicates the 95% confidence interval.

To estimate the growth advantage provided by chrysolaminarin in natural conditions, we considered six diatom blooms reported in the literature^73–78^ (Fig. 6a, colored circles), spanning different locations and seasons. For each bloom, we determined the diel cycle and light intensity at the depth of the bloom using data collected by the Moderate-Resolution Imaging Spectroradiometer (MODIS) of NASA’s Aqua satellite (Supplementary Methods). We then ran simulations with our model using the extracted bloom conditions and calculated the growth advantage. This analysis revealed that, for conditions typical of natural blooms, utilizing chrysolaminarin at night provides a considerable growth advantage, ranging from 8.8% to 14.4% for the six blooms considered (Fig. 6b, colored circles).

To assess how this growth advantage can influence diatom biomass production in the ocean, we used our model (with and without storage) to simulate a 4-day^76–80^ bloom for each of the six environmental bloom conditions^73–78^ (Fig. 6c). We found that, by the end of the four days, diatoms using chrysolaminarin for growth in darkness reach a 2.0- to 3.3-fold larger population than diatoms not using storage. This represents the upper limit of the growth advantage, as it assumes no losses due to grazing or slower division rates due to nutrient limitation, both of which are expected to decrease the actual realized advantage in the environment. After 10 days, the difference grows to 7.9- to 16.8-fold (Supplementary Methods). These predictions suggest that, while the direct benefit of using chrysolaminarin to fuel growth in darkness applies to a relatively small fraction of cells in a population, the cumulative effect over the course of a multi-day bloom is considerable and applies to conditions that are typical of natural blooms in the ocean across different seasons and geographic locations.

Finally, since day/night cycles and light intensities in the environment are closely linked to changes in seasonality, we used the model to estimate the growth advantage provided by heritable chrysolaminarin stores across the year. For the same six reported bloom locations, we extracted the average diel cycle and light intensity for each month of the year 2023. Then, for each bloom location and month, we ran a 4-day growth simulation and extracted the growth advantage at the end of the 4-day period (Fig. 6d). Model results indicate that the growth advantage of heritable chrysolaminarin stores is maximal in spring and autumn conditions and lower in winter and summer conditions. In winter, cells would benefit most from chrysolaminarin, but because the light intensity is very low, very little chrysolaminarin can be accumulated during the brief light periods. Thus, given the lower levels of chrysolaminarin, the realized growth advantage in winter is predicted to be lower. In summer, high light intensities and long days would favor chrysolaminarin accumulation, but its importance for growth is smaller because dark phases are short. In spring and autumn, instead, dark phases are long enough for heritable energy reserves to have a significant impact on growth, while daylight intensities are sufficient to allow substantial chrysolaminarin accumulation.

## Discussion

Population growth rate, a key parameter determining the role of phytoplankton in oceanic primary production^81,82^, ultimately results from the growth and division cycles of individual cells. Yet, the fact that we know surprisingly little about the growth dynamics of individual cells limits our ability to understand and ultimately predict how populations grow and respond to environmental change. This is particularly relevant for diatoms exhibiting asynchronous growth and division^5–16^. Due to this asynchrony, traditional bulk growth assays cannot quantify the generation times of individual cells. At the same time, single-cell measurements using light^83,84^ or holographic microscopy^85^ have to date reported generation times of maximally 215 cells^84^, owing partially to the challenge of maintaining many cells in focus over prolonged times^84^ and to accurately quantify their growth using image analysis^83^. Such throughput is too low to resolve the fine-grained, hourly effects of diel cycles on *T. pseudonana*’s generation times that our work uncovered.

Here we introduced a microfluidics-based approach to track the growth of thousands of individual, asynchronously growing *T. pseudonana* cells. Analyzing the growth trajectories of individual cells relative to their time of birth in the diel cycle revealed that cells born after sunset arrested growth until sunrise, whereas those born before sunset continued to grow through the night using energy previously stored as chrysolaminarin. Remarkably, if non-arrested cells enter darkness with sufficient chrysolaminarin storage, they attain generation times that are nearly as short as those of cells growing entirely in light. Lipid accumulation and consumption have also been shown to be coupled to the diel cycle in the diatom *Phaeodactylum tricornutum*^86^, suggesting that lipids can also serve as diel energy reserves. In *T. pseudonana*, while we did observe a modest (1.5x) increase in lipids from sunrise to sunset (Extended Data Fig. 9a), the increase in chrysolaminarin (10.8x) was considerably more pronounced (Extended Data Fig. 9b), pointing to chrysolaminarin as the main diel energy reserve in T*. pseudonana*.

Chrysolaminarin plays a key role in the marine carbon cycle, serving as a diel energy reserve in diatoms, as previously recognized in both laboratory^63,66^ and environmental studies^62,87^. Yet, the growth advantage afforded by chrysolaminarin reserves for asynchronous diatom populations had remained elusive. Our findings suggest that the amount of chrysolaminarin stored is important for non-arrested cells, enabling them to grow rapidly even when most of their cell cycle occurs in darkness. Ultimately, this mechanism allows non-arrested cells to decouple their growth from light availability.

Our results show that chrysolaminarin accumulation depends on the light history not only of the cells themselves, but also of their parents. In diatoms, chrysolaminarin is stored in a large vacuole localized within the cytoplasm^70,88,89^. Contrary to other organelles such as chloroplasts and mitochondria^90^, little is known about the inheritance of vacuoles in diatoms during cell division. Yet, in other eukaryotic organisms such as yeast^91^ and plants^92^, daughter cells can inherit parts of their parental vacuoles. We therefore propose that chrysolaminarin in *T. pseudonana* could be inherited from one generation to the next through vacuole inheritance from parent to daughter cell, passing on to newly formed cells part of the energy accumulated by their parents. Previous work^84^ demonstrated that hypothecal-derived daughter cells divide faster than epithecal-derived ones in the diatom *Ditylum brightwellii.* We speculate that, if vacuoles (and therefore chrysolaminarin) are asymmetrically partitioned during mitosis, this could generate such asymmetry in division rates among sister cells, especially for those growing mostly during darkness.

The different division patterns (synchronous vs. asynchronous) among phytoplankton groups might reflect divergent evolutionary strategies. Asynchronously growing diatoms are often considered successful r-selected species, producing many offspring when conditions are favourable. In contrast, more synchronously growing phytoplankton groups, such as Prochlorococcus, Synechococcus and dinoflagellates, tend to be more K-selected, generating fewer offspring. Given that diatoms often exhibit higher growth rates compared to other phytoplankton groups^20,21,27,28^, we speculate that the utilization of chrysolaminarin as diel energy storage, by enhancing asynchronous growth, might have evolved as a beneficial strategy for diatoms in their ecological niche, contributing to their high growth rates under favourable conditions.

Fast asynchronous growth equips *T. pseudonana* with an ecological advantage under various environmental conditions. We have shown that cells can complete a division cycle with only minimal light during asynchronous growth, using stored chrysolaminarin during darkness, thus avoiding energy constraints that otherwise slow growth during the night. Our model results show that heritable diel energy reserves boost asynchronous population growth rates, leading to a 2.0-to 3.3-fold increase in population size over a 4-day bloom period across different environmental bloom scenarios. Thus, if the mechanism of heritable energy reserves sustaining growth in darkness is widespread among diatoms, we expect it to be a major contributing factor to the fast, asynchronous population growth rates observed across diatoms during bloom scenarios, when cells are actively growing in the euphotic zone under abundant light and nutrient conditions. In contrast, when cells leave the sunlit regions, for example through vertical mixing or sinking, other traits will become more important, such as the ability to respond to long-term starvation.

Ultimately, it is the interplay of many traits—ranging from diel energy storage to protective silica frustules and resilience to starvation—that underpins the ecological success of diatoms in contemporary oceans.

Our model does not account for trade-offs for chrysolaminarin storage, thus representing a maximal possible growth benefit. If cells did not store fixed carbon as chrysolaminarin, this carbon could be allocated towards immediate biomass synthesis, biosynthetic machinery (e.g. ribosomes)^93–96^ or secretion of dissolved organic matter into the phycosphere^79^. Diverting fixed carbon into storage instead of biomass-producing machinery can slow down overall growth by reducing the rate at which new biomass can be synthesized. At the same time, our results show that stored carbon can increase growth during the night. Thus, a balance might exist between investing carbon into biosynthetic machinery and storing carbon to upkeep high biomass synthesis rates at night. Diatoms likely have optimized this partitioning to maximize overall population growth rate. Regarding immediate biomass synthesis, the trade-off with storage will depend on what nutrient limits growth. Phytoplankton growth is often limited by nitrogen^97–99^ silicate^76,100,101^ or iron^98,102,103^, though in some scenarios it is limited by inorganic carbon^104^.

When inorganic carbon is limiting, the carbon allocation trade-off between biosynthesis and storage will likely be more important than if other nutrients are limiting. Finally, the trade-off with secretion into the phycosphere is more difficult to evaluate. Diatoms can exude a significant fraction of fixed carbon (e.g., 20% of total fixed carbon^79^): this can stimulate metabolic interactions with bacteria^105^, which can benefit diatom growth by providing limiting nutrients such as vitamin B_12_^106,107^, yet the relative growth advantage compared to storing carbon for growth at night might be very context dependent.

Previous work has demonstrated that during rapid, nutrient-replete growth, cells allocate more fixed carbon to nitrogen-rich biosynthetic machinery (e.g., ribosomes) and less to carbon-rich storage compounds such as chrysolaminarin, resulting in a higher N:C (Nitrogen:Carbon) ratio compared to slower, nutrient-limited growth^93,94^. Ecosystem models have shown that such shifts in carbon allocation across macromolecules can explain global patterns in elemental stoichiometry (N:C:P) and the nutrient-dependent acclimation of phytoplankton in the ocean^108,109^. At the same time, our findings demonstrate that storage compounds are important during nutrient-replete conditions by sustaining nighttime growth. To reconcile these results, we argue that the flux of chrysolaminarin across the diel cycle is an important parameter in explaining the relative importance of storage compounds during rapid growth. During nutrient-replete growth, the flux of chrysolaminarin is high, with chrysolaminarin contributing from 7% (sunrise) to 40% (sunset) of total biomass in *P. tricornutum*^110,111^. In contrast, under nutrient limitation, the diel chrysolaminarin flux is strongly reduced and chrysolaminarin can accumulate to constant levels^112^. Therefore, the average chrysolaminarin concentration across the diel cycle during rapid growth might be lower (owing to the higher flux) than in nutrient-limited conditions, resulting in lower N:C ratios. Overall, during rapid growth, cells invest more in proteins and RNA^93,94^ and at the same time a higher flux of chrysolaminarin reduces the average concentration of chrysolaminarin across the entire diel cycle, both of which contribute to higher N:C ratios. During slow growth, cells invest less into proteins and RNA and at the same time chrysolaminarin builds up (as it can no longer be used to fuel growth), both of which contribute to lower N:C ratios. Thus, we argue that the flux of storage compounds should be considered when assessing the relative importance of chrysolaminarin during rapid growth. Indeed, analysis of surface ocean stoichiometry in the Atlantic and Indian oceans revealed significant diel variation in the N:C ratio, declining during the day and rising during the night, with the amplitude of the N:C ratio variation positively correlating with growth rate^113^.

Diatoms, although globally distributed, are most prominent at high latitudes, and undergo strong seasonal cycles, forming large-scale blooms typically during spring^114–118^. Diatoms are thought to dominate these spring blooms partially due to their high growth rates^114,119,120^. The predictions of our model, based on six historical diatom bloom locations with prevalence of high-latitude regions, indicate that the growth advantage afforded by heritable energy reserves at these high latitude regions is maximal during spring and autumn months. We propose that under those conditions dark phases are still long enough for heritable energy reserves to play an important role, while at the same time light intensities during the day are high enough to allow for substantial chrysolaminarin accumulation. As such, the ability to use chrysolaminarin to support growth at the intermediate diel dark periods found in spring might contribute to the prevalence of diatoms at high latitudes, boosting their population growth rate during spring blooms compared to other phytoplankton groups.

How widespread the growth advantage provided by heritable diel energy reserves is among diatom species remains an open question. Diatoms are remarkably diverse^118^ and some diatom species such as *T. weissflogii*^64,121,122^ might have two (or more) light-dependent cell-cycle checkpoints, which would reduce the ability of heritable diel energy reserves to increase population growth, as multiple checkpoints (especially those located at the end of the cell cycle) would increase the fraction of cells which dark arrest during the night, leading to an overall reduction in population growth rate. Yet, chrysolaminarin has been detected in many diatom taxa in the laboratory^88^ and our metagenomics analysis (Extended Data Fig. 7a) revealed that some of the globally most abundant diatom genera such as *Chaetoceros* and *Thalassiosira*^118^ likely possess the necessary enzyme to synthesize chrysolaminarin, indicating the widespread importance of chrysolaminarin in bloom conditions across diatoms in the ocean. In this context, our experimental approach and individual-based model provide a blueprint to study the role of heritable energy reserves across other species, which will ultimately be important to determine how broadly utilized this ecological strategy is in the ocean.

Finally, our results demonstrate that measuring phytoplankton growth at the single-cell level can reveal important new aspects of the growth dynamics of populations and ultimately, we expect, furthering our understanding of phytoplankton physiology, ecology and impact on global primary production. Therefore, we envision that the single-cell approach to phytoplankton growth introduced here will be broadly valuable in studying phytoplankton physiology and ecology, also in the context of environmental perturbations and interactions with other microbial players such as heterotrophic bacteria.

## Methods

### Diatom medium and batch culturing

Non-axenic cultures of *T. pseudonana* (CCMP 3367) were obtained from the National Center for Marine Algae and Microbiota (NCMA) at Bigelow NCMA and grown in 25 cm² cell culture flasks (Nunc EasYFlask, Thermo Scientific) using L1 medium (Bigelow, NCMA). A sea salt mix (Instant Ocean) was used to achieve 3.5% salinity in the L1 media. The cultures were maintained in a phytochamber (AlgaeTron AG230, Photon Systems Instruments) at 22 °C with a 14 h light / 10 h dark cycle under white LED light (100 μmol photons m^−2^ s^−1^). Cultures were maintained in semi-continuous mode by diluting cultures every 14 days 1:30 with fresh L1 medium under a laminar flow hood. To measure population growth of batch cultures, three replicate cultures were grown under a 14 h light / 10 h dark diel cycle and sampled daily for 14 days, starting immediately after splitting the replicate cultures. Cells were counted using a CytoFLEX S flow cytometer (Beckman Coulter). *T. pseudonana* cells were identified using forward scatter (FSC) and side scatter (SSC) (Extended Data Fig. 5a).

### Microfluidic device design and fabrication

To conduct the single-cell growth experiments, a microfluidic device with one serpentine shaped channel (32 cm length × 800 µm width × 200 µm height) was designed. Microfluidic devices were fabricated using standard soft lithography techniques. SU-8 photoresist (KAYAKU Advanced Materials) was deposited on a silicon wafer via photolithography to prepare the microchannel master mold, which was then silanized with trichloromethylsilane (Sigma-Aldrich, CAS Nr. 75-79-6) before use. Polydimethylsiloxane was mixed with 10% wt/wt cross-linking agent (Sylgard 184 Silicone Elastomer Kit, Dow Corning) and cast onto the master mold. The devices were subsequently cured at 80 °C for at least 12 h. Following fabrication, PDMS devices were bonded to a glass slide (Avantor) by plasma treating each interacting surface for 1 min. The assembled chip was baked at 80 °C for 2 h and cooled overnight before use.

### Microfluidics experimental procedure

Cells were grown to early exponential phase in batch cultures (Extended Data Fig. 1a). During microfluidic experiments, cells were exposed to one of three different diel regimes: short day with 6 h light / 18 h dark, long day with 14 h light / 10 h dark, or continuous day with 24 h light. Prior to experiments, batch cultures were pre-exposed to the light regime for one complete diel cycle (24 h). On the day of the experiment, cells were diluted to a concentration of 5×10^4^ cells/ml in fresh L1 medium. The cells were loaded into the microfluidic device and the device was sealed before being mounted on an inverted microscope (NIKON Ti2 Eclipse). To provide actinic illumination during the day, a low-angle ring light (LA-120W, Smart View) was mounted around the microfluidic device (Extended Data Fig. 1b) and connected to a digital timer (Brennenstuhl). Light intensity of the ring light was set to either 100 μmol photons m^−2^ s^−1^ or 43 μmol photons m^−2^ s^−1^ depending on the experimental condition using a light meter (WALZ ULM 500) with a spherical sensor. A full-case incubator (Life Science Instruments) maintained the temperature around the microscope and microfluidic device at 22 °C to match the temperature of the phytochamber. Cells were allowed to settle for 30 min prior to starting image acquisition.

### Microscopy image acquisition

Imaging was performed on a Nikon Eclipse Ti2 Eclipse inverted microscope with an automated stage controller (Prior Scientific) and laser-based autofocus system (PFS, Nikon). The imaging light intensity, supplied by a CoolLED pE-100, was set to 15% of maximum intensity (corresponding to 8 μmol photons m^−2^ s^−1^ on the microfluidic device) with an exposure time of 30 ms and passed through a neutral density filter. Bright-field time-lapse images were acquired using 20× magnification at 30–50 positions along the microfluidic channel at a frequency of 1 frame every 15 min for a total duration of 72 h (‘long day’ and ‘short day’ conditions) or 52 h (‘continuous day’ condition). Since cells showed displacement in the *z*-direction during the experiment, a 5 × 7 μm *z*-stack (z-direction = height direction of microfluidic device) was recorded at each position using a sCMOS camera (Hamamatsu Orca Flash, 6.5 μm per pixel). The image acquisition procedure was automated using Nikon Elements.

### Image processing

Raw images were processed with an image analysis pipeline using custom python, ilastik ^123^ and Fiji ^124^ scripts. First, for each position and time point, the 5 × 7 μm *z*-stacks were converted into a single image using a minimum intensity projection, followed by median-background subtraction across all time points using a rolling ball algorithm to reduce background noise. Next, images were segmented by separating pixels into foreground (cells) and background with a random forest classifier using the pixel classification workflow of ilastik. The resulting pixel probability maps were converted into binary images containing cells and background. Touching objects were separated using the watershed function in Fiji. Cells were tracked and division events recorded using the Linear Assignment Problem (LAP) tracking algorithm of the trackmate ^125^ plugin for Fiji. Tracking results were processed using a custom python script to assemble lineages and compute single-cell growth time series, which start with the birth of a new cell and end with its division into two new cells. The generation time is defined as the time between birth and division of a cell. Objects not corresponding to cells, non-growing cells and mistakes during automated tracking were excluded using manually defined thresholds for minimal (30 μm^3^) and maximal (1000 μm^3^) size, minimal growth rate (slope of linear fit of size vs. time > 0), and minimum track length (18 frames). These thresholds were applied automatically during image processing.

### Datasets and statistical analysis

Data were analyzed using Python or Graphpad Prism (10.1.2, GraphPad Software). Since time of both birth and division need to be recorded during the experiment in order to calculate generation times, only cells born within the first 48 h of a 72 h experiment (long day and short day) or 35 h of a 48 h experiment (continuous day) were included in the analysis (Extended Data Fig. 2b).

Including cells born close to the end of the experiment would lead to a bias towards short generation times, because cells born close to the end of the experiment but with long generation times would not divide within the course of the experiment and so would not be captured within the data. In all figures, statistical tests are two-tailed, *n* refers to the number of single cells measured and *N* to the number of independent replicates is stated. The significance level for all statistical tests was set at p = 0.05.

### Tracking cell volume doubling during division cycles

Tracking individual growth trajectories revealed that cells continuously increase in volume through their division cycle (Extended Data Fig. 8c), doubling in volume from birth to division (Extended Data Fig. 8a,b). The generation time refers to the time a cell takes to complete a full division cycle, during which a cell grows and ultimately divides. Given that biomass derived from volume measurements correlates strongly with experimentally measured biomass in single phytoplankton cells^85^, we assume that the doubling in volume we observe corresponds to a doubling in biomass, as expected during balanced exponential growth. Further, no significant difference exists in either birth volume (M.W.U test, *p* = 0.14) or division volume (M.W.U test, *p* = 0.97) between the two diel cycles.

### Measurement and analysis of chrysolaminarin and lipids

To measure chrysolaminarin levels during the diel cycle, cells were grown in batch culture and either sampled at 2 h intervals or at sunset only. Cells were fixed using 10% Lugol’s solution (Sigma) and stained for 10 min using 1mg/ml aniline blue (Thermo Scientific). Samples were analyzed using a CytoFLEX S flow cytometer (Beckman Coulter). Samples were measured at flow rate of 60 ul/min for 1 min, resulting in minimally 8252 and maximally 9702 cells measured per sample. Cells were identified by gating an unstained population in forward scatter (FSC) and side scatter (SSC) (Extended Data Fig. 5a). Aniline blue fluorescence was recorded using the KO-525 channel. Chrysolaminarin staining was confirmed independently for a separate test sample using fluorescence microscopy by exciting stained cells using 1% power of a 395 nm laser (Spectra X) and recording the emission using a DAPI filter cube (Chroma Technology). To measure lipids, cells were stained for 10 min using 2 µg/ml BODIPY (Thermo Scientific).

Samples were measured using the same flow cytometer settings and gating strategy as for chrysolaminarin. BODIPY fluorescence was recorded using the FITC channel.

### Calculation of total photons received

To estimate the total photons (in μmol) received by a cell from birth to division, we derived the average surface area (in m^2^) of a cell from birth to division and multiplied it with the light intensity (100 or 43 μmol photons m^−2^ s^−1^) experienced by the cell and the cell’s generation time (in s). Using a cell’s average radius in µm, and approximating a cell as a sphere, the surface area of a cell is determined by the sphere surface area formula A = 4πr^2^. The surface area in µm² is then converted to m². While this represents the upper limit of total photons received by a cell (not all photons will be absorbed by the photosystems of a cell), the subsequent analysis does not rely on absolute quantification of photons but the relative difference between conditions (Long day 100 m^−2^ s^−1^ vs. Long day 43 μmol m^−2^ s^−1^ vs. Short day 100 m^−2^ s^−1^)^126^

### Mathematical model of single-cell diel growth dynamics

A detailed description of the model can be found in the Supplementary Methods

### Detection of Chrysolaminarin Synthase gene in Tara oceans dataset

A search for the *T. pseudonana* chrysolaminarin synthase gene homologues was performed in the *Tara* oceans eukaryotic Metagenome-Assembled-Genomes (MAGs) and Single-cell Assembled Genomes (SAGs) datasets. The BLASTN algorithm based search was performed using the *T. pseudonana* chrysolaminarin synthase nucleotide sequence (THAPSDRAFT_12695, NCBI ID: 7450323) with an e-value ≤1e-10 as threshold. The analysis was conducted through the Ocean Gene Atlas v2.0 (https://tara-oceans.mio.osupytheas.fr/ocean-gene-atlas/) ^126^ and the resulting taxonomic distribution chart was downloaded.

## Data availability

All tracking results and simulation outputs required to reproduce the findings of this manuscript and examples of raw microscopy data are available in a pre-publication dataset at Dryad: https://datadryad.org/share/2jH0YXj3DMTHuGBIY0UvaI1HqjNqNQWVTueNHLAHMOk.

Full raw microscopy data (∼10TB) will be available at publication or if requested during peer-review. Tara Oceans data are available at: https://tara-oceans.mio.osupytheas.fr/ocean-gene-atlas/

## Code availability

All analysis scripts, code and mathematical models required to reproduce the findings of this manuscript are available in a pre-publication dataset at Dryad: https://datadryad.org/share/2jH0YXj3DMTHuGBIY0UvaI1HqjNqNQWVTueNHLAHMOk.

## Acknowledgments

We thank R. Naisbit for help with editing the manuscript, J. Słomka and R. Naudascher for discussions and E. Burmeister for supporting the laboratory work. We gratefully acknowledge funding from a Gordon and Betty Moore Foundation Symbiosis in Aquatic Systems Initiative Investigator Award (GBMF9197; https://doi.org/10.37807/GBMF9197), the Simons Foundation through the Principles of Microbial Ecosystems (PriME) collaboration (grant 542395FY22), Swiss National Science Foundation Sinergia grant CRSII5-186422, and the NCCR Microbiomes, a National Centre of Competence in Research funded by the Swiss National Science Foundation (grant numbers 51NF40_180575 and 51NF40_ 225148) to R.S.; from a H2020 MSCA Individual Fellowship (886198), and the European Union and a Bavarian Gender Equality Grant (2024) from the University of Bayreuth to C.M.P.

## Author contributions

CRediT authorship contribution statement

Conceptualization: OM, DAB, FC, CMP, JMK, RS

Methodology: OM, TR, DAB, RS

Investigation: OM

Formal analysis: OM, TR

Visualization: OM, JMK

Funding acquisition: RS

Project administration: OM, RS

Supervision: CMP, JMK, RS

Writing – original draft: OM, JMK

Writing – review & editing: OM, TR, DAB, CMP, JMK, RS

## Competing interests

All authors declare that they have no competing interests.

## Materials & correspondence

Correspondence should be addressed to Oliver Müller: and Roman Stocker

---

**Extended Data Fig. 1.**
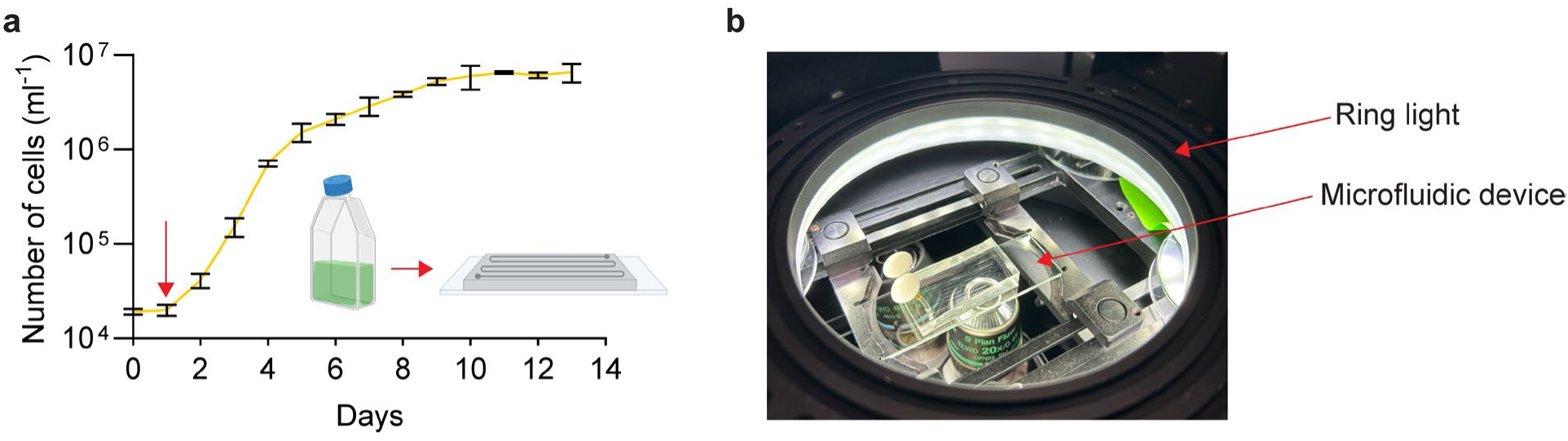
Microfluidic set-up to track diatom growth at single-cell resolution. **(a)** Bulk population growth curve of *T. pseudonana*. Exponential phase starts at day 1 (red arrow) and lasts until day 4. Prior to microfluidic experiments, cells were grown to early exponential phase (red arrow, day 1) then loaded into a microfluidic device to conduct single-cell growth experiments. Yellow line represents the mean, black error bars the standard deviation of *N* = 3 independent replicates. **(b)** The experimental microfluidic set-up, with the microfluidic device in the center, mounted on a Nikon Eclipse Ti-2 microscope. The ring light provides even actinic illumination during the experiment.

**Extended Data Fig. 2.**
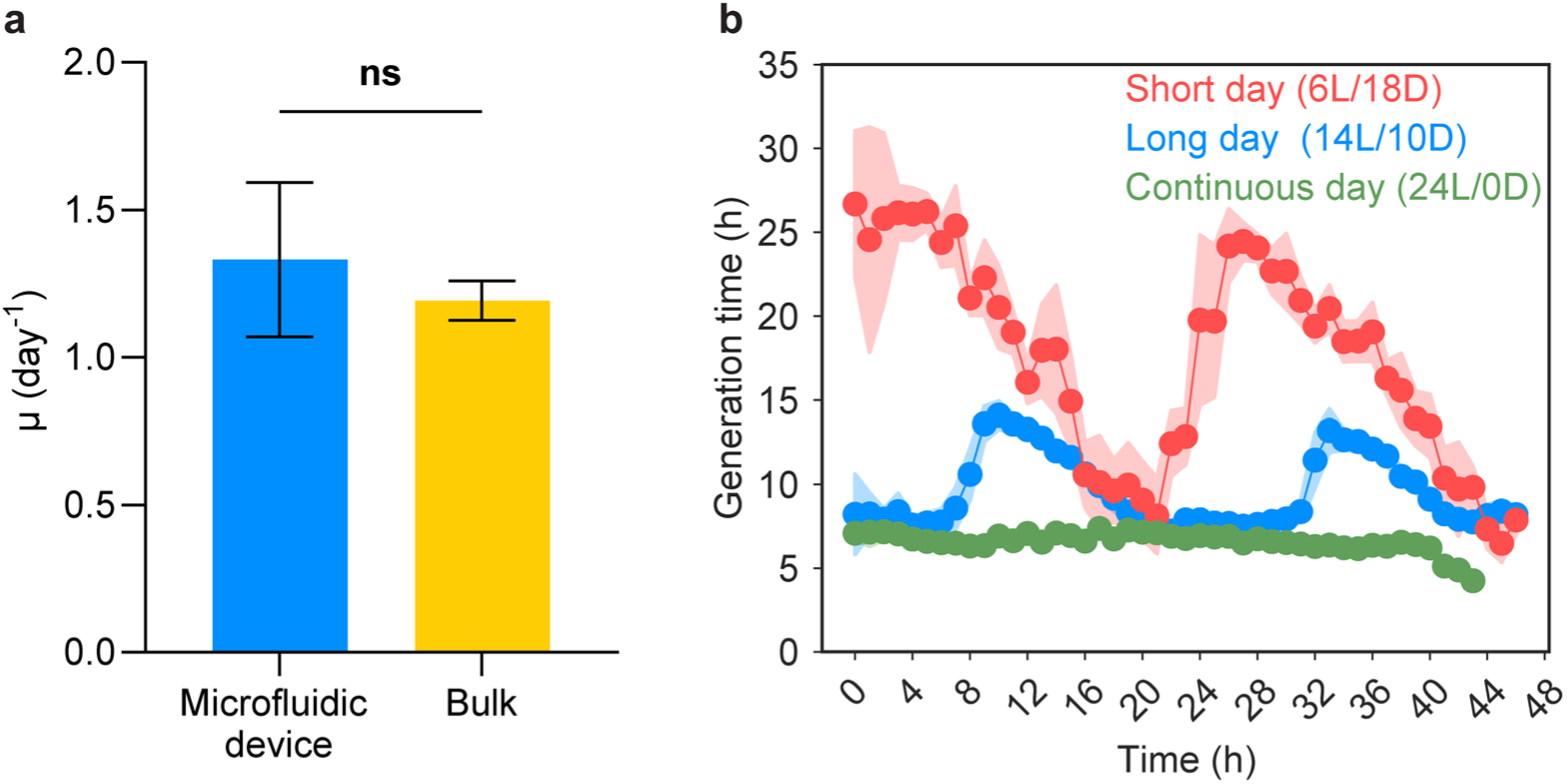
The microfluidic device enables reliable extraction of generation times in *T. pseudonana.* **(a)** Mean population growth rate of cells growing in a 14 h light / 10 h dark diel cycle estimated from cells growing in a microfluidic device (blue) or in standard batch culture (yellow). Population growth rate in the microfluidic chip was estimated by counting the number of cells (N_0_) at the beginning (t_0_) and end (N_1_) of the experiment after 3 days (t_1_) within the microfluidic chip and calculating the daily population growth rate μ = ln (N_0_ / N_1_) / (t_0_ – t_1_). Using the same formula, μ was calculated for the bulk culture (data from Extended Data Fig. 1a) during the exponential phase (between day 1 and day 4). Bar plot represents the means, error bars the 95% CI. The estimates of population growth rate did not differ between microfluidic chip and bulk culture (*p* = 0.52, M.W.U. test, *N* = 3 independent replicates for microfluidic chip, *N* = 3 independent replicates for bulk cultures). **(b)** Generation time of cells as a function of the time point of the experiment at birth, for cells growing under conditions of continuous day (green), long day (blue) or short day (red). Instead of combining cells form two days into one diel cycle (Fig. 2b) cells are plotted across the entire duration of the experiment. Points show the mean of 1-hour bins, and shading the 95% confidence interval. Generation time shows oscillation during the experiment due to diel cycles, but these oscillations stay constant for the 48 h of the experiment, indicating that there is no slowdown of growth rate due to nutrient limitation or other factors.

**Extended Data Fig. 3.**
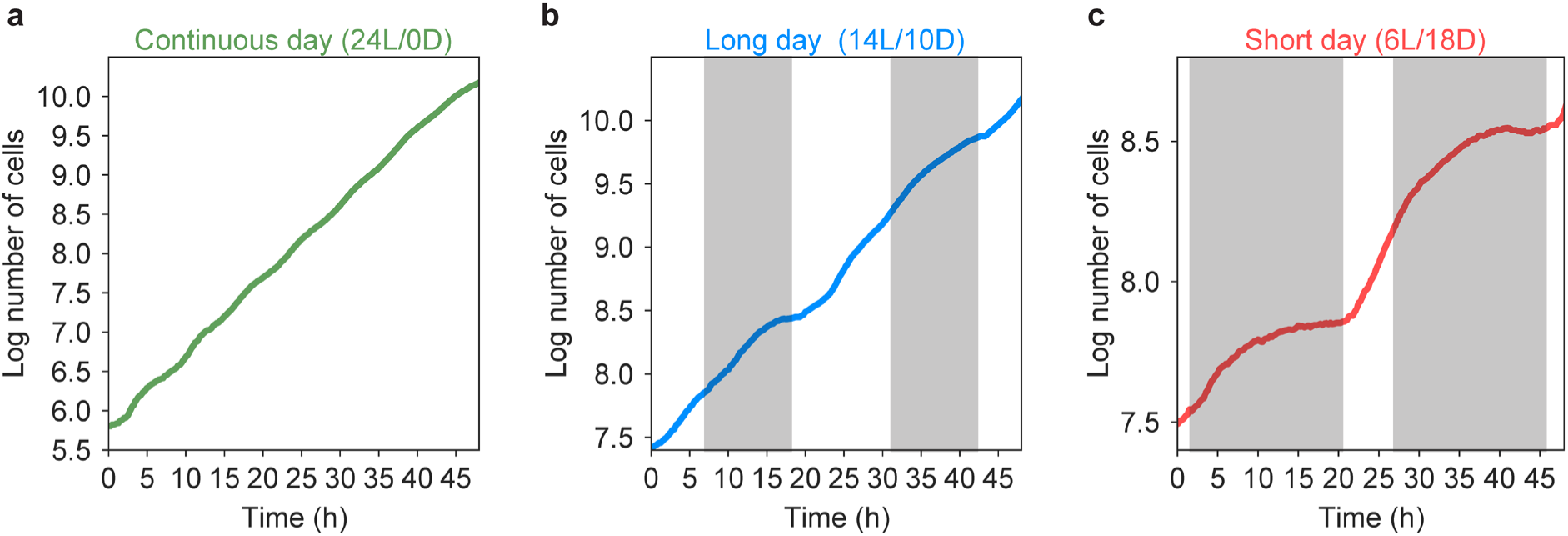
Cultures of *T. pseudonana* grow asynchronously, with divisions occurring throughout the diel cycle. Logarithm of cell count within the microfluidic chip as a function of the time point within the experiment for conditions of **(a)** continuous day, **(b)** long day, and **(c)** short day. Grey shaded areas in (b) and (c) indicate the dark phase.

**Extended Data Fig. 4.**
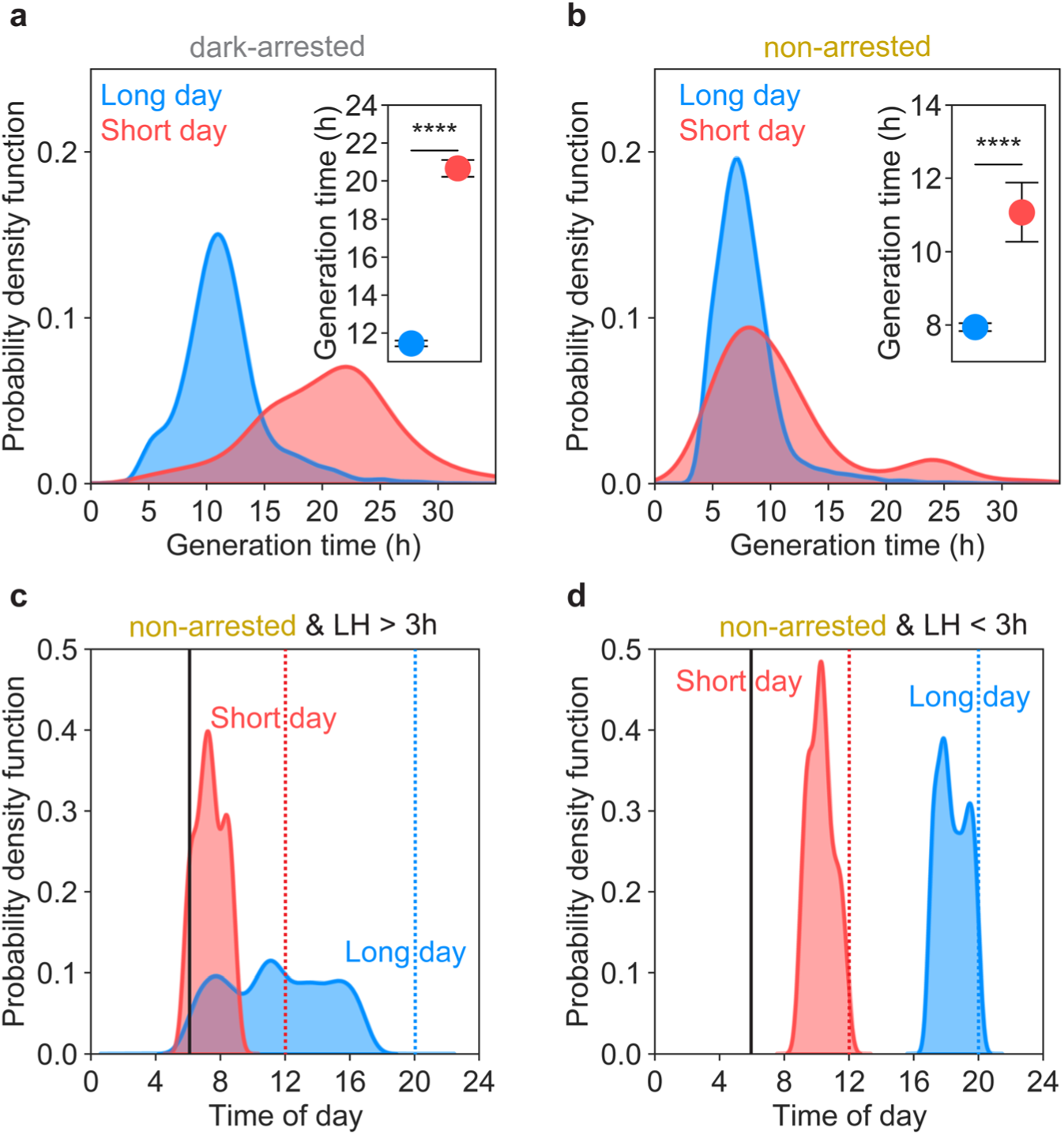
Dark-arrested and non-arrested cells grow faster in the long-day compared to the short-day diel cycle. **(a)** Generation time distributions for dark-arrested and **(b)** non-arrested cells for long-day (blue) and short-day (red) conditions. Inset shows the mean generation time and 95% CI for long and short day. In the long day, dark-arrested (*p* <0.0001, M.W.U. test, *n_l_*_ong day_ = 2206 cells, *n*_short day_ = 799 cells) and non-arrested (*p* <0.0001, M.W.U. test, *n_l_*_ong day_ = 3249 cells, *n*_short day_ = 255 cells) cells grow significantly faster than in the short day. **(c)** Distributions of time of day at birth for non-arrested cells that received more than 3 h of light (*LH* > 3) in the long-day (blue) and short-day (red) conditions. Vertical black line indicates sunrise, vertical dashed lines indicate sunset. Cells with *LH* > 3 in the short day originate from early in the light period, cells in the long day originate from the early and middle parts of the light period. **(d)** Distributions of time of day at birth for non-arrested cells that received less than 3 h of light (*LH* < 3) in the long-day (blue) and short-day (red) conditions. Vertical lines as in (c). Cells with *LH* < 3 in both the long and short day originate from the end of the light period.

**Extended Data Fig. 5.**
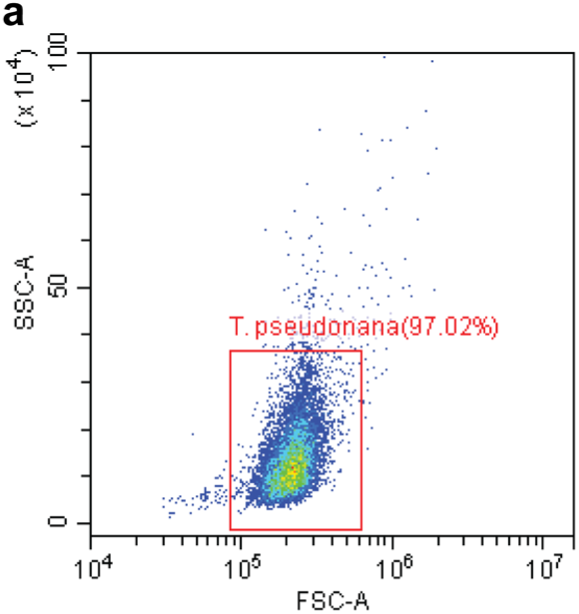
Gating strategy used to collect flow cytometry data. **(a)** T. pseudonana population was characterized according to side scatter (SSC) and forward scatter (FSC).

**Extended Data Fig. 6.**
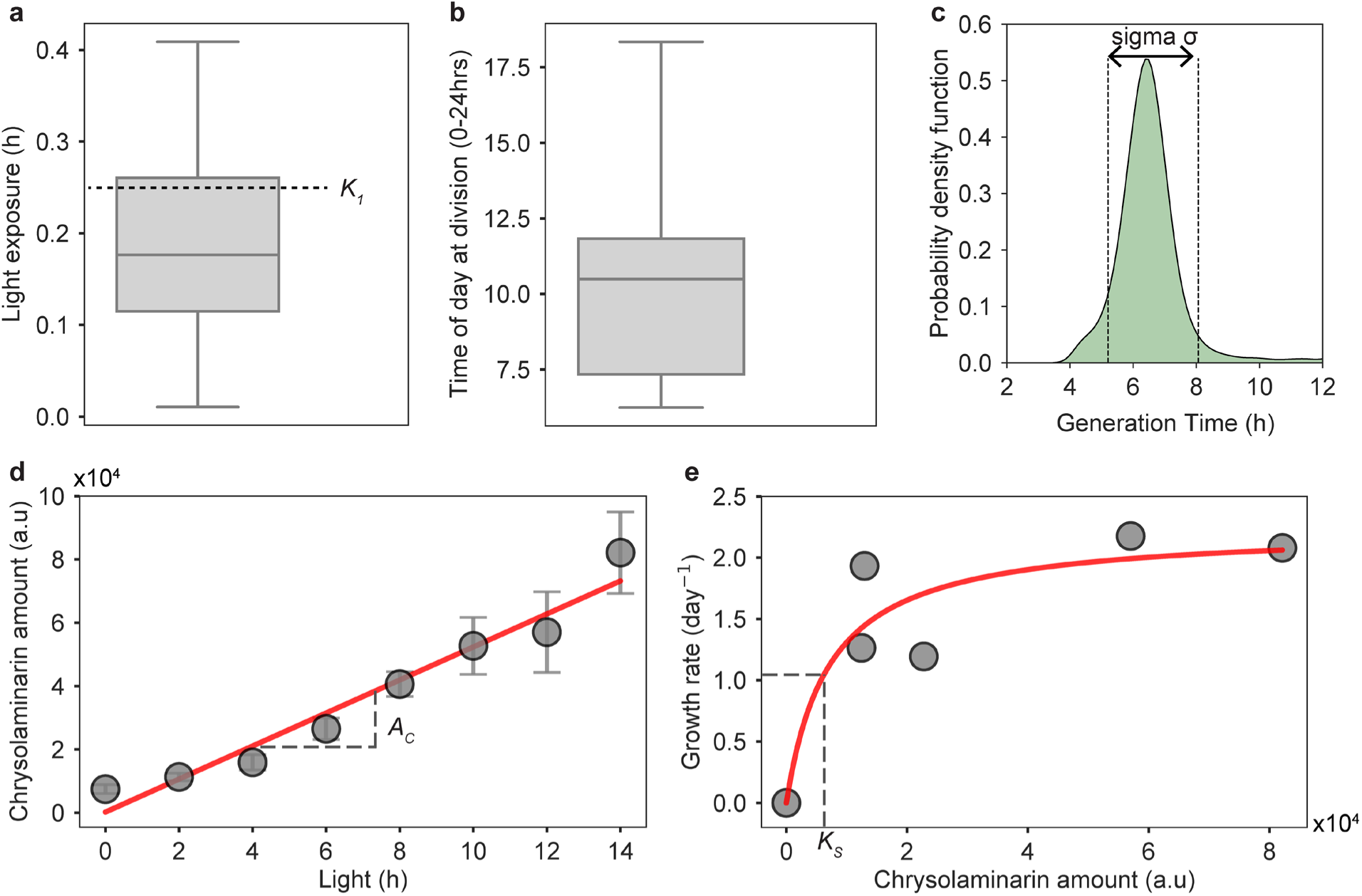
Extraction of model parameters from experimental data. **(a)** Estimation of the light-dependent cell cycle segment K_1_. Light exposure for cells that receive the least amount (bottom 1%) of light in the long-day condition. **(b)** Estimation of the light-independent cell cycle segment N_2_. Time of division for dark-arrested cells born between 20:00 and 21:00 in the long day condition. In (a) and (b), the line indicates the median, box boundaries the 25^th^ and 75^th^ percentile, and error bars the range. **(c)** Estimation of sigma σ. Generation time distribution of cells growing in continuous illumination (green). Dashed lines indicate +/-standard deviation of generation times around the mean generation time. Sigma corresponds to two times the standard deviation. **(d)** Estimation of chrysolaminarin accumulation rate A_c_. Increase of chrysolaminarin with duration of the light period during the long-day condition. Data is the same as in Fig. 4b. Grey points represent the average chrysolaminarin signal; red line is the linear fit. Grey dashed lines indicate slope of linear fit which corresponds to A_c_ (5212 chrysolaminarin fluorescence/hour). **(e)** Estimation of Half-saturation constant K_s_. Monod equation (red curve) is fitted to chrysolaminarin and the corresponding growth rates obtained from experiments in long and short-day condition (grey points). Point (0/0) was not experimentally measured but is based on the assumption that when no storage compounds are present, cells cannot produce energy or carbon skeletons for growth to fuel divisions in darkness. Chrysolaminarin data same as in Fig. 4b. Growth rate data same as in Fig. 2. Grey dashed line indicates K_s_. For (e) and (e), the chrysolaminarin signal is fluorescence intensity × 10^4^.

**Extended Data Fig. 7.**
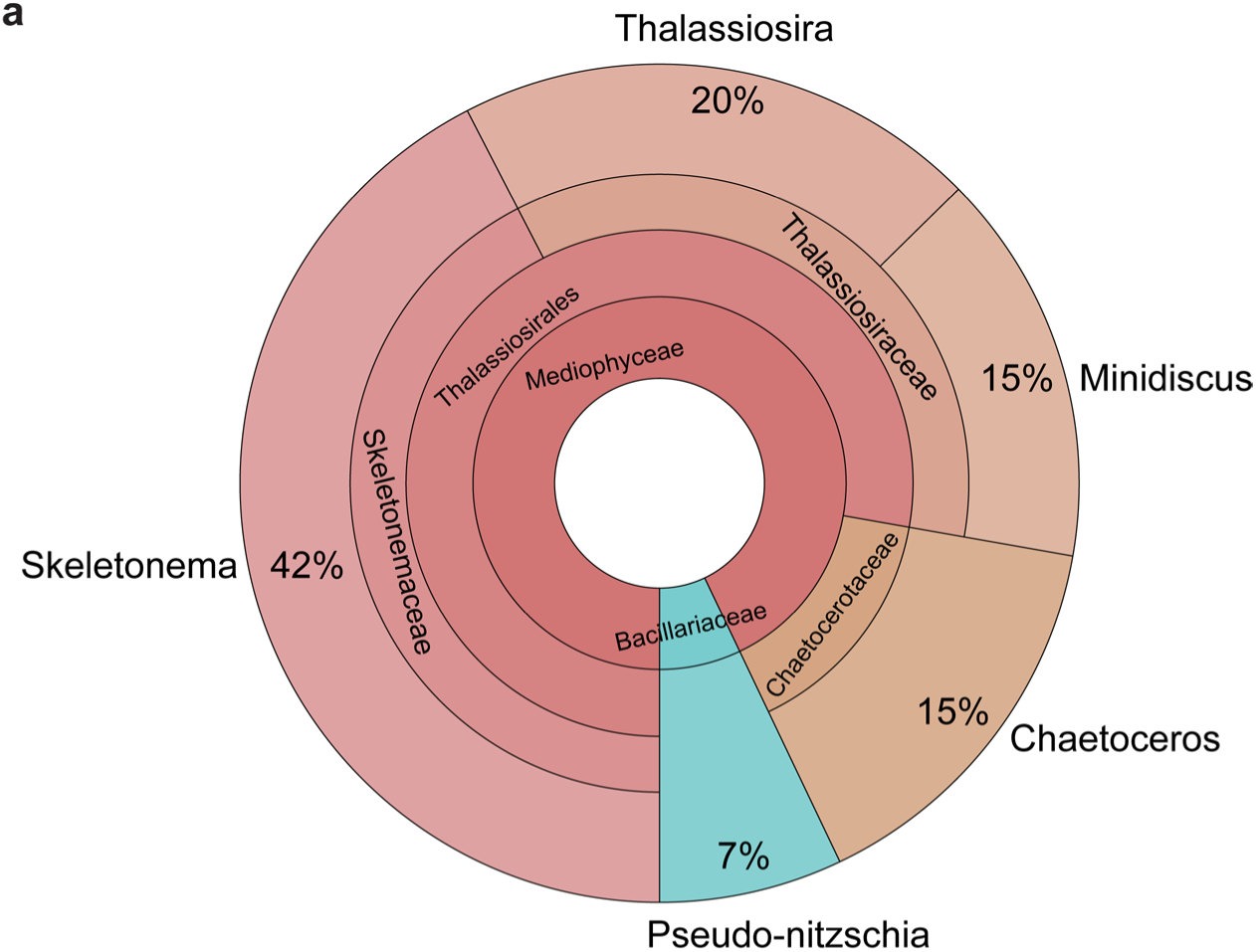
Chrysolaminarin synthase taxonomic distribution in the ocean. **(a)** Taxonomic distribution of the *T. pseudonana* chrysolaminarin synthase sequence homologues in the eukaryotic Metagenome-Assembled-Genomes (MAGs) and Single-cell Assembled Genomes (SAGs) *Tara* Oceans dataset.

**Extended Data Fig. 8.**
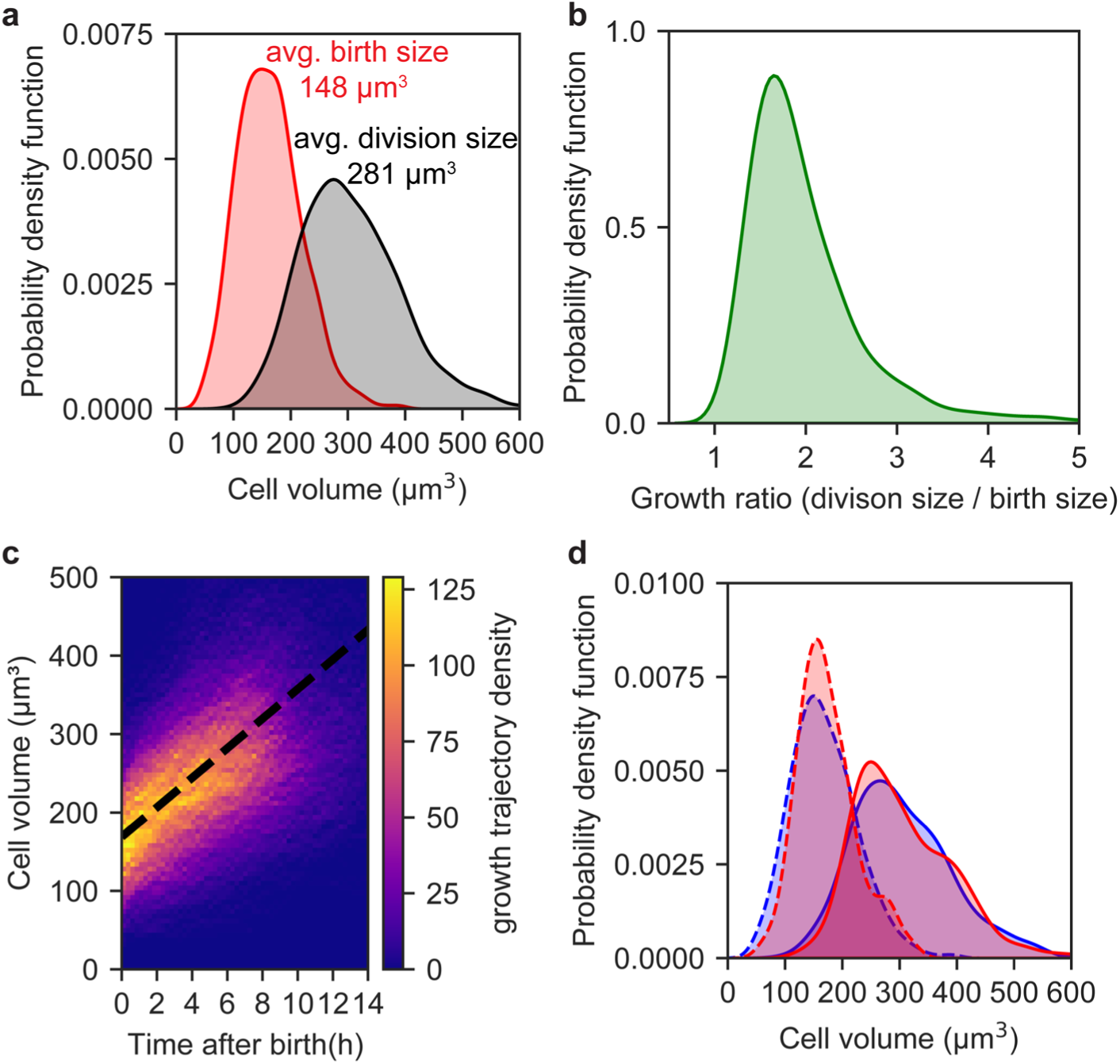
Single-cell growth dynamics of non-arrested cells with LH < 3. **(a)** Probability density function of cell volume at birth (red) and division (black). **(b)** Probability density function of volume increase of single cells across a complete division cycle. **(c)** Heatmap of single-cell volume trajectories as a function of time after birth. Each trajectory corresponds to the volume evolution of an individual cell from birth until division, aligned at time zero (birth). The color intensity reflects the density of trajectories at each volume–time coordinate, with warmer colors (yellow) indicating a higher number of overlapping trajectories and cooler colors (blue) indicating fewer. The linear fit across all trajectories (black dashed line) has a slope of 18.9 µm^3^/hour. **(d)** Probability density function of cell volume at birth (dashed) and at division (solid) in long (blue) and short (red) day. No significant difference in either birth volume (M.W.U test, *p* = 0.14) or division volume (M.W.U test, *p* = 0.97) between the two diel cycles.

**Extended Data Fig. 9.**
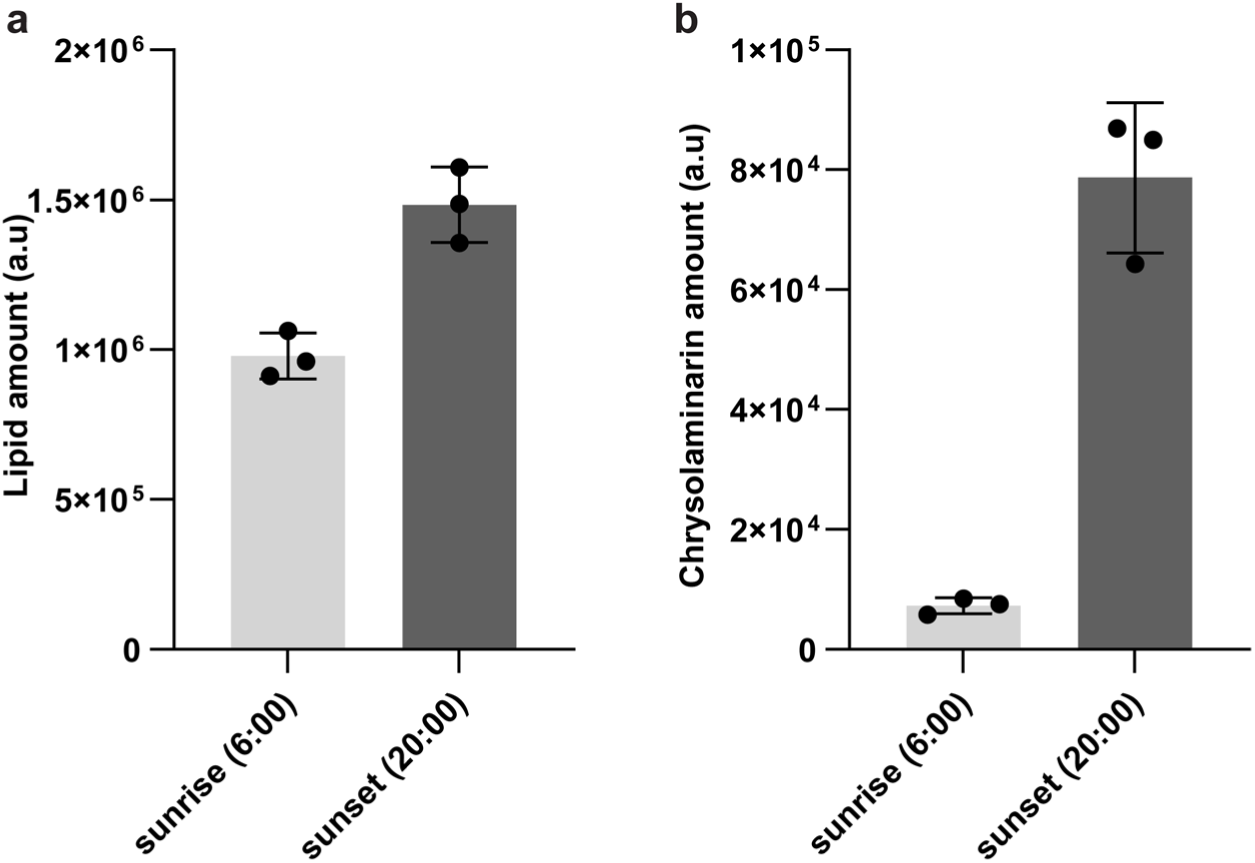
Diel patterns of chrysolaminarin and lipid storage. **(a)** Lipid and **(b)** chrysolaminarin accumulation across the light period in the long day condition. Cells increased (a) lipids 1.5-fold and (b) chrysolaminarin 10.8-fold from sunrise to sunset. Bar plots indicate the mean, and error bars the standard deviation of *N* = 3 independent replicates.

**Extended Data Fig. 10.**
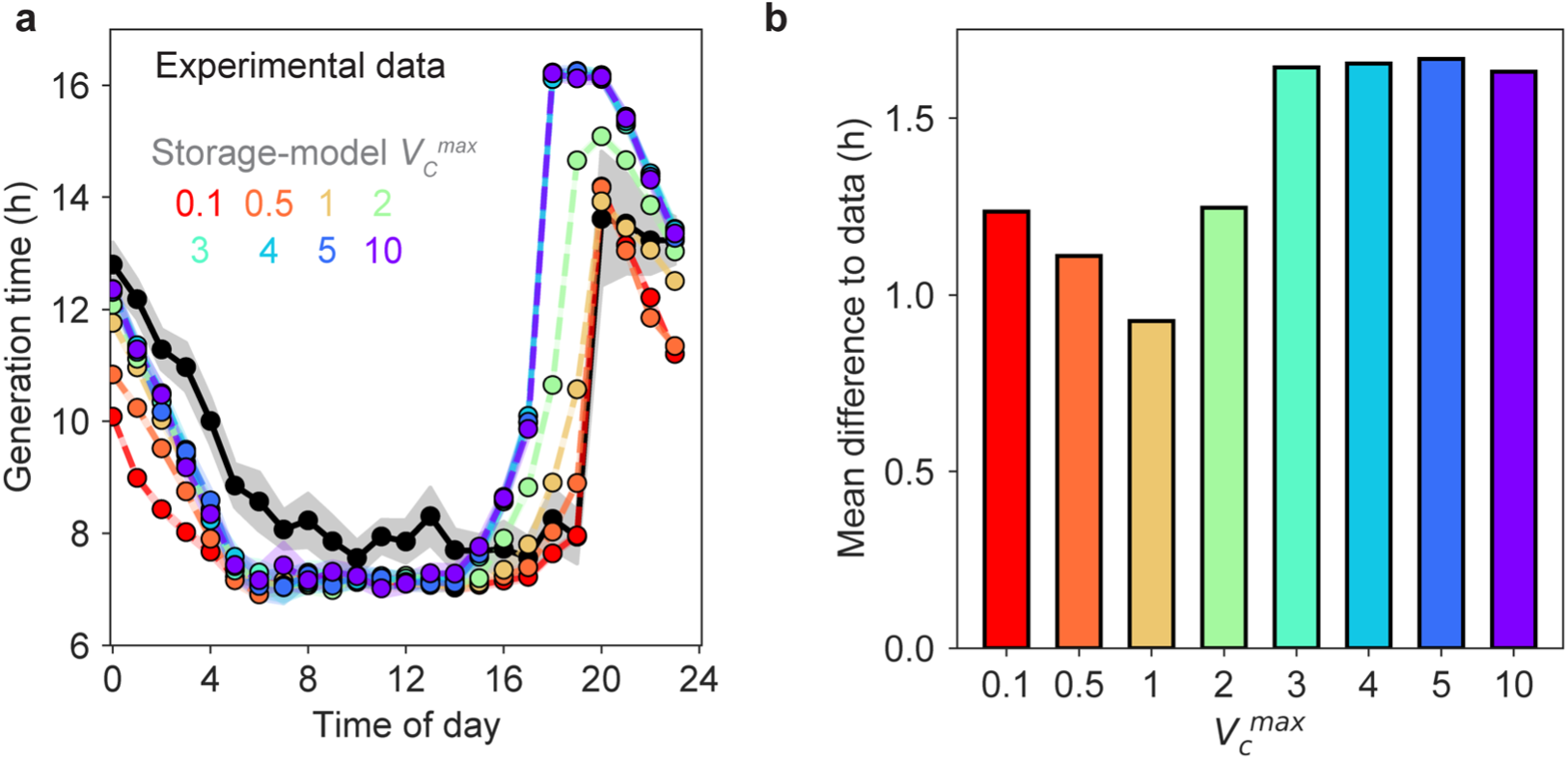
**(a)** Generation time of cells as a function of the time of day when the cell was born for cells growing in a long day. Solid black line is the experimental data (from Fig. 2b) for the long day. Colored dashed lines are the predictions of our Storage model for the long day for different values of *V_C_^max^* (0.1 – 10). Circles show the mean of 1-hour windows and shading indicates the 95% confidence interval. **(b)** Mean generation time difference between predictions of the Storage model for different values of *V_C_^max^* (same color as in (a)) compared to the experimental data.

## Supplementary Information

### Supplementary Text

#### An overview of diatom growth dynamics in light/dark cycles

The patterns of diatom population growth in diel cycles have been studied for decades, yet only recently have molecular studies started to unveil some of the mechanistic cellular processes which ultimately give rise to these growth patterns. This section aims at putting old and new population-based studies in context with the molecular mechanisms we now know to be important in shaping diatom growth in diel cycles, as well as highlighting how the single-cell approach introduced in this manuscript can shed new light on diel diatom growth dynamics, revealing processes in asynchronously growing diatom populations normally hidden by population-based growth studies.

#### Asynchronous diatom population growth under non-limiting conditions

Traditionally, diel growth patterns of diatoms and other phytoplankton have been studied at the population level by measuring the increase in cell number across the diel cycle, resulting in classic population growth curves. Synchronous divisions occur when all cells within a population divide during one or sometimes two defined time windows of the diel cycle and, importantly, the population division rate is zero outside of these windows^13^. In some of the literature^13^, the distinction is made between synchronous division (all cells divide each day during a narrow time window) and phased division (only a proportion of the cells divide each day). Importantly, in both scenarios, divisions occur only during a restricted time window, with the population division rate being 0 outside that window. Thus, in our manuscript we capture both patterns under the term ‘synchronous division’.

In contrast, in asynchronous populations there is always a fraction of the population dividing at any given point in the diel cycle and the population division rate is always greater than zero^13^. Overall, diatoms tend to divide asynchronously with respect to the diel cycle under non-limiting conditions, with many studied species showing population-level division rates greater than zero across the entire diel cycle^5–16^. This means that a fraction of the population is dividing at any given time point in the diel cycle, leading to divisions occurring during both day and night.

When discussing asynchronous population growth, we must also acknowledge that the degree of asynchrony can vary among species and with environmental conditions. Some species show a very high degree of asynchrony, which is characterized by a near-constant population division rate across the entire diel cycle, leading to a near-constant increase in cell number during both day and night. An example of a highly asynchronous growing species is *T. pseudonana* ^6,10,13^ (Extended Data Fig. 3B) which shows near-constant population division rates in diel cycles when the dark phase is moderately short or short. Other diatom species such as *T. weissflogii*, while still growing asynchronously with division rates always greater than zero, show multiple peaks with increased division rates across the diel cycle^13,14^. Experiments where the same species were grown in different light/dark cycles have also revealed that the degree of asynchrony at the population level is influenced by the ratio of the light/dark phases^13^ (Extended Data Fig. 3 A-C).

Thus, while diatoms grow asynchronously under non-limiting conditions, the degree of asynchrony can vary both among species and depend on the light/dark ratio of the diel cycle.

#### Molecular mechanisms coupling diatom cell division to diel cycles

To understand how phylogeny and environment (e.g. different light/dark cycles) can shift the degree of asynchrony in diatoms, one must consider the molecular mechanisms at play which ultimately control cell division in diel cycles. Light-dependent cell cycle checkpoints have emerged as a key determinant of diatom division patterns in diel cycles^29–31^. Light-dependent cell cycle checkpoints require cells to be exposed to light to pass through a checkpoint within their cell cycle and ultimately divide. A light-dependent cell cycle checkpoint located in G1 has been identified and characterized at the molecular level in *P. tricornutum*^29,30^. Other diatoms, such as *T. weissflogii*, have a second light-dependent checkpoint located in G2/M^64,121,122^. Genomic studies and our data support the existence of a single light-dependent checkpoint located early in the cell cycle in *T. pseudonana* (see Discussion).

The number of light-dependent checkpoints and their location within the cell cycle will, in combination with the diel regime, shape the observed population division pattern. To illustrate this, let us consider the simplest case where cells are placed in prolonged darkness (e.g. for 24 h, see section “Synchronization protocols“). In prolonged darkness, all cells will eventually arrest at a light-dependent cell cycle checkpoint. After being released back into the light, a species with one checkpoint will then divide synchronously with one single peak, because the entire population arrests at the same point in the cell cycle. In a species with two checkpoints, a fraction of the population will arrest at each checkpoint. When released into light, there will be two characteristic peaks because some cells need more time to divide (those arrested earlier in their cycle) than others (those arrested later in their cycle). Of course, once cells are released from prolonged darkness into continuous light or a diel light/dark cycle, they will rapidly desynchronize due to the inherent variability in division time among cells, which is not linked to the diel cycle. This inherent cell-to-cell variability of generation times is captured in our model by the parameter sigma. The larger this variability, the faster the population will eventually desynchronize again upon re-illumination.

The circadian clock represents another mechanism by which diatom physiology is coupled with the diel light/dark cycle^33,34^. Although the circadian clock regulates numerous physiological processes in diatoms, the extent to which cell division is governed by the circadian clock versus light-dependent cell cycle checkpoints is still an open question. In our experiments with *T. pseudonana*, we did not observe division patterns under continuous light that would suggest circadian control. However, recent studies in *P. tricornutum* have proposed a potential role for the circadian clock in modulating division timing^33,34^. It is therefore plausible that light-dependent checkpoints and circadian regulation act together to coordinate cell division within the context of diel light/dark cycles of some diatom species.

#### Influence of different light/dark ratios on observed asynchrony at the population level

Given current understanding of light-dependent cell cycle checkpoints, it is not surprising that population-level division patterns vary with the light/dark ratio of the diel cycle. For instance, when the dark phase is relatively short or moderate, *T. pseudonana* maintains a nearly constant population division rate throughout both light and dark periods. This occurs because the generation time of non-arrested cells (e.g., ∼8 h for *T. pseudonana* under long-day conditions) is comparable to the duration of the dark phase (e.g., ∼10 h in our long-day experiments). Since *T. pseudonana* requires light only at the onset of its division cycle, cells born shortly before sunset will complete their division just before the sunrise. At the population level, this results in divisions occurring throughout the entire dark period. However, when the dark phase is extended, as in our short-day condition, the fraction of arrested cells increases, and the overall population division rate declines during the night. This trend is evident at the population level in our data (Extended Data Fig. 3), where under short-day conditions division rates progressively decrease through the night, reaching a minimum just before sunrise. Yet, while coarse-grained assessments of e.g. population division rates are possible using average population-based studies, how rapidly the underlying cells really grow and divide is impossible to assess with traditional population-scale approaches.

#### Influence of nutrient limitation on observed asynchrony at the population level

Besides light-dependent checkpoints, nutrient limitation can also arrest the cell cycle of diatoms. For example, different diatom species starved of silica will arrest their cell cycle at different locations of their cell cycle^64^. Upon re-exposure to the limiting nutrient, the population will undergo a few rounds of division in a more synchronized manner before de-synchronizing again.

#### Synchronization protocols

To study cell-cycle-related processes using traditional, bulk, population-based approaches, it is essential that most cells progress through the cell cycle stages simultaneously. Otherwise, averaging signals from cells at different stages obscures stage-specific patterns^6^. Because diatoms grow asynchronously under nutrient-replete conditions, this has prompted researchers to use both light and nutrient starvation to try and transiently synchronize diatom populations.

For instance, studies that identified genes involved in light-dependent cell cycle checkpoints required the population to transition synchronously through the cell cycle^29,30^. Sampling an asynchronous population under non-limiting conditions for e.g. transcriptomic analysis would blur stage-specific gene expression profiles, making it impossible to determine which genes are active in G1, S, or G2/M phases. Because diatoms are naturally asynchronous under non-limiting conditions, researchers have relied on methods to transiently synchronize diatom populations prior to taking measurements.

One such approach is to use prolonged (24 h) darkness to arrest all cells within a population. After release back into the light, the majority of the population undergoes a few rounds of synchronized division. This approach has been used for example to study the molecular mechanism of light-dependent cell cycle checkpoints^29,30^.

Alternatively, silica starvation has been used to arrest cells at the silica-dependent cell cycle checkpoint, and upon re-introducing silica into the growth medium, study the synchronized formation of new silica frustules^6^. Again, without this synchronization, studying for example which genes are involved in silica deposition would be difficult, as cells in an asynchronously growing population do not undergo new cell wall formation at the same time.

#### Limitations of population-based studies of asynchronously growing diatoms

While population-based studies of diel growth dynamics in diatoms can reveal broad patterns, such as whether division occurs continuously or in distinct peaks, they cannot resolve the division timing of individual cells. Although population-level data indicate when divisions occur, the inherent asynchrony makes it impossible to determine how fast individual cells grow or what their specific light histories are. Consequently, in asynchronous populations, metrics such as the average population growth rate or instantaneous division rate provide little insight into single-cell behaviour. This limitation is made evident by our data: for instance, under short-day conditions, the calculated average generation time is not representative of what individual cells experience, as most cells divide either much faster or much slower than this mean value. To accurately quantify how individual cells grow within an asynchronous population, a single-cell tracking approach such as the one presented in this study is required. This approach enables measurement of each cell’s generation time and allows one to relate growth dynamics to the light history of the cell and of its progenitors. Such tracking of single cells along lineages made it possible in this study to reveal how cells in an asynchronously growing population can maintain rapid growth during darkness and to identify the underlying mechanism: heritable diel energy reserves of chrysolaminarin, passed on from parent cells growing during the day to daughters growing during the night.

### Supplementary Methods

#### Short description of the individual-based model of *T. pseudonana* growth

The aim of the model is to capture how chrysolaminarin storage affects the growth dynamics of individual diatom cells and, consequently, the population growth rates under diel light/dark cycles. Diatom cells must complete both light-dependent and light-independent processes during their cell cycle to progress past a critical transition point (called light dependent cell cycle checkpoint, *T*) and divide. Light-dependent cell cycle checkpoints have been previously identified for the diatom cell cycle^61^. Our single-cell growth data is consistent with such a light dependent checkpoint located early in the cell cycle in *T. pseudonana* (Fig. 3b and main text). To complete the cell cycle, progress through the checkpoint, and ultimately divide, the cell must meet a required level of light exposure. Previous models^71,72^ have incorporated such a transition point by assuming that the cell cycle temporarily arrests at the transition point during darkness when the light exposure during the previous light period was insufficient to meet the cell’s light requirements. As such, the cell cycle arrests at the light-dependent transition point until the next light period, where cells can continue to fulfil their light requirements and eventually cross the transition point. Such models assumed that the rate of cell growth (i.e., the speed of cell cycle completion) was constant in time and solely determined by the presence or absence of light as a binary input. As such, cells who have passed the checkpoint grow at a constant rate in both light and darkness. However, our experimental data shows that for cells growing mostly in darkness, the rate of growth is not constant, but depends on the light history of the parent cells, which determines the amount of stored energy in the form of chrysolaminarin. Therefore, to quantify the growth advantage of these heritable diel energy reserves, improving on the existing models was required.

Here, we expand on the previous models by adding the experimentally observed dependency of cell growth on stored chrysolaminarin to the existing framework of light dependent cell cycle checkpoints. While cells use the energy provided by light to grow during the day, they rely on stored energy (chrysolaminarin) to grow during darkness. The growth in darkness depends directly on the intracellular quantity of stored chrysolaminarin. Specifically, we observed that the growth rate of cells growing mostly in darkness displayed a Monod relationship with respect to the amount of stored chrysolaminarin (Extended Data Fig. 6e). Therefore, we model the cells growth rate in darkness using Monod kinetics, with stored chrysolaminarin serving as a limiting nutrient. As such, the difference to previous models is that, while in previous models cells grow at a constant rate during darkness, in our model, this rate is directly dependent on stored chrysolaminarin. This approach better captures the experimentally observed generation times of cells which grow mostly during darkness (Fig. 5b,c and main text) and enables us to quantify the growth advantage provided by chrysolaminarin at the single-cell and population level. We implemented an individual-based model where we simulate the progression of individual cells through their cell cycle. We tracked the time of birth and the time of division for each cell and all of its progeny growing in diel light/dark cycles. From these simulations of individual cell growth, population metrics such as growth rate or number of cells can easily be derived at any given time point.

#### General model description and formalism

Here, we present the details of our model of diatom growth dynamics in diel light/dark cycles. We implemented an individual based approach where the cell’s growth dynamics during darkness is dependent on its chrysolaminarin storage. Further, in our model, chrysolaminarin is accumulated during the day and passed on from parent to daughter cells so that a cell’s chrysolaminarin storage depends upon the light history of its progenitor cells.

The diatom cell cycle consists of the eukaryotic cell cycle segments G1 (Gap 1, cell growth and preparation for DNA replication) / S (Synthesis, replication of DNA / G2 (Gap 2, preparation for cell division / M (Mitosis, separation of duplicated DNA and cell division). The progression from one segment to the next is safeguarded by cell cycle checkpoints, ensuring favourable conditions before progressing with the next segment. In diatoms, progression through G1 and G2 can be dependent on light^61^. Therefore, light-dependent cell cycle checkpoints in G1 and G2 integrate environmental cues into the cell cycle progression of diatoms.

In our model, assuming spatial homogeneity, all cells are subjected to homogeneous illumination during diel light/dark cycles. Based on our experimental data and previous reports of light dependent checkpoints in *T.pseudonana*^64^, we assume the existence of one light-dependent checkpoint located early in G1 (see Fig. 3b and main text). As in previous models^71,72^, the cell cycle is simplified from G1/S/G2/M phases down to two sets of light-independent processes and one light-dependent process to achieve division (Fig. 5a). We define *k_1_* as the minimum number of hours of light exposure required for a cell to complete a division cycle (Fig. 5a). The time intervals required to complete light-independent processes are measured in hours and respectively denoted as *n_1_* and *n_2_* (Fig. 5a). The cell must first complete the initial segment of light-independent processes, which last for *n_1_* hours, and be exposed to light for a minimum duration of *k_1_* hours to pass the checkpoint *T*. The second set of light-independent processes, which last *n_2_* hours, begins immediately after crossing the checkpoint. We define the age of a cell using two variables, *n* and *k*, which, respectively, represent the cell’s progression, i.e. hours elapsed, in light-independent and light-dependent processes^71^. During the day, when the cell is exposed to light, each hour, the cell undergoes one hour of ageing, both in light-dependent and in light-independent processes. During darkness, cell growth rate is dependent on the internal chrysolaminarin concentration, as explained in the next section.

#### Chrysolaminarin accumulation dynamics and cells growth speed

We characterize the cell’s age through two distinct variables, one accounting for light dependent age, *k*, and one accounting for light-independent age, *n*. We assume light dependent processes to progress at a constant rate over time

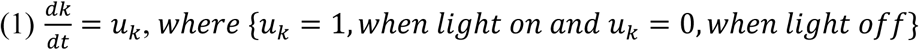

In our experiments, we observed that during the light phase, cells maintain a constant growth rate (continuous day and long day, Fig. 2b) while storing chrysolaminarin (Fig. 4b) and that the concentration of chrysolaminarin within the cells increases linearly in time (Fig. 4b and Extended Data Fig. 6d). In contrast, we observed that cells growing mostly during darkness (non-arrested cells) slow down when they have little chrysolaminarin stored (short day at 100 µmol m^-2^ s^-1^ and long day 43 µmol m^-2^ s^-1^) but maintain rapid growth when more chrysolaminarin is stored (long day 100 µmol m^-2^ s^-1^ and short day 233 µmol m^-2^ s^-1^). These observations led us to hypothesize that the cell’s aging process, for light independent processes, progresses linearly with time in the presence of light and is based on chrysolaminarin concentration *C_chrysolaminarin_* consumption during darkness. We incorporate the dynamics of *C_chrysolaminarin_* in our model. During the light phase, the concentration increases linearly with time

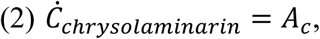

where *A*_*c*_ is the accumulation rate, measured experimentally for *T. pseudonana* cells. Since we did not observe a saturation of the chrysolaminarin signal during the light period of the tested diel cycles (long day and short day, Fig. 4b), we assume no saturation of the chrysolaminarin storage capacity in our model. Next, during darkness, the consumption of chrysolaminarin follows a Monod kinetics (during light, there is no consumption of chrysolaminarin)

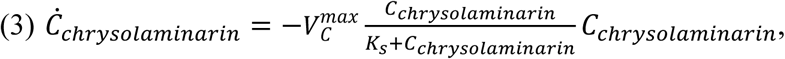

where *K*_s_ represents the half-velocity Monod constant, determined experimentally. The parameter *V_C_^max^* represents the maximum biomass production rate. *V_C_^max^* has the units of hours^-1^.

Since we did not have a priori knowledge of *V_C_^max^*, we ran our simulations for the Storage model with different values for VCmax (Extended Data Fig. 10) and found that the model aligns best with the experimental data when *V_C_^max^* = 1 hours^-1^. We therefore chose *V_C_^max^* = 1 for our simulations. As in previous models, cell growth in light is constant (equation (4), first line) and during the dark phase is limited to cells that have completed the light-dependent processes and therefore have passed the light-dependent cell cycle checkpoint during the previous light phase (equation (4), third line). In previous models, growth during darkness is always constant (*dn/dt = 1* when light is off). Since this does not accurately capture the experimentally observed growth dynamics of *T. pseudonana* cells growing mostly in darkness, we sought out to expand on these models by adding the observed growth dependency on stored chrysolaminarin. Therefore, our model makes the growth during darkness dependent on stored chrysolaminarin (equation (4)). Cells grow during dark phases by consuming their stored chrysolaminarin, following Monod kinetics. The aging of light-independent processes is modelled as

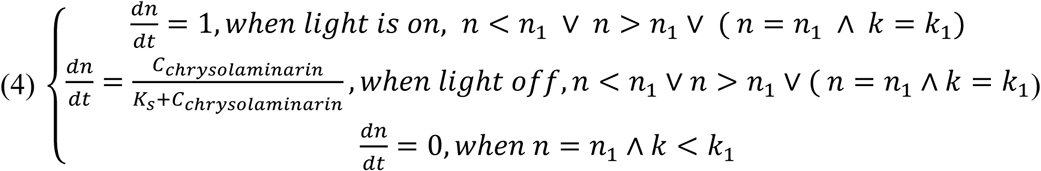

Here we assumed *n*_1_to be the minimum time devoted to processes happening before transition T, *n*_2_the minimum time between transition *T* and division, and *k*_1_ the minimal light requirements. The cells’ aging halts at the checkpoint *T* if either *n_1_* or *k_1_* processes are not satisfied. Cell doubling occurs when both light-dependent and light-independent processes are satisfied. Overall, the minimum time required for a cell to double is therefore *t_m_*

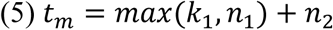

##### Population growth

To numerically replicate our experimental setup, we simulate the growth of a population of cells exposed to diel light/dark cycles. Cell cycle phases are not strictly deterministic, but instead display a certain amount of variability^71^. Following established approaches^71,72^, and to maintain a simple model formulation, we assume that cell parameters within the population are distributed according to

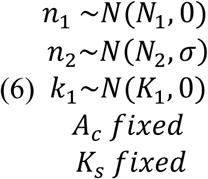

Where we set the entire variability only in the cell-cycle segment happening after the checkpoint. Indeed, to account for the intrinsic biological variability of cells, the cell parameter *n_2_* follows a normal distribution with mean *N*_2_and standard deviation *σ*. The variability *σ* represents the inherent biological differences between cells within the population. We estimated such variability fitting a gaussian distribution to our experimental data and fixing *σ* as twice the standard deviation of the experimentally observed division times. Each cell’s age evolves according to the dynamics in Eq. (1) and Eq. (4), until division. Upon division, cell C1 splits into two newborn cells C2 and C3, which age parameters are reset to zero *C*2(*n*, *k*) = (0,0), *C*3(*n*, *k*) = (0,0). The chrysolaminarin storage of the dividing cell is equally split amongst newborn cells 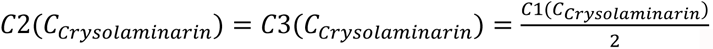. Through this process, cells continuously pass on their stored chrysolaminarin to their progeny, enabling cells which receive little light themselves, but inherit lots of chrysolaminarin from their parent cell, to maintain fast growth in light-independent segments during darkness.

Cell division occurs always at *k* = *k*_1_. The duration of light-independent processes within a cell is assumed to follow a normal distribution. Specifically, the probability *p*(*n*) that a cell will divide between light-independent ages *n* and *n* + *dn* is

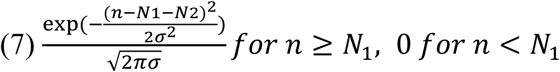

We list the units of all the model parameters in the Supplementary Table 1 below:

**Supplementary Tabel 1:** Description of model parameters.

| Parameter | Description | Units |
| --- | --- | --- |
| $C_{Chrysolaminarin}$ | Chrysolaminarin concentration | Fluorescence (arbitrary units) |
| $A_C$ | Chrysolaminarin accumulation rate | Fluorescence (arbitrary units) / hour |
| $K_S$ | Monod half-velocity constant | Fluorescence (arbitrary units) |
| $N_1$ | Light independent segment 1 | Hours |
| $N_2$ | Light independent segment 2 | Hours |
| $K_1$ | Light dependent segment 1 | Hours |
| $\sigma$ | Variability in $N_2$ | Hours |
| $V_C^{max}$ | Yield | Hours<sup>-1</sup> |

##### Simulations

We generate a population of cells where *n* and *k* age variables are initialized following an arbitrary initial distribution. The model parameters *K_1_*, *N_1_*, *N_2_*, *A_C_*, *K_S_* and σ were estimated for *T. pseudonana* using the experimental data presented in this article (see section Parameter estimation from data; Extended Data Fig. 6). When describing the parameters in general, we refer to them in lower case letters. When we refer to the average values that these parameters take in our specific case of *T. pseudonana*, we refer to them in upper case letters. Cells are entertained for 5 diel dark/light phases before performing measurements of population averaged quantities such as average population growth rate and average population generation time.

##### Parameter estimation from data

We estimated the model parameters *K_1_*, *N_1_*, *N_2_*, *A_C_*, *K_S_* and σ for *T. pseudonana* using the data presented in this article.

*K_1_*: *K_1_* represents the number of hours of light exposure required for a cell to complete the light-dependent processes before crossing the transition point *T*. *K_1_* can be determined from single-cell growth data by examining the light exposure of the non-arrested cells with the least light exposure in the entire dataset. This represents the minimal light exposure necessary for a cell to divide. We identified these cells in our dataset, sorting them by the amount of light they received, and selecting the 1% of cells experiencing the shortest light exposure. The mean light exposure for these cells was 0.18 hours (Extended Data Fig. 6a). Since our model is discretized in 0.5-hour intervals, we defined *K_1_* as 0.5 hours.

*N_2_*: *N_2_* represents the amount of time required for light-independent processes after the transition point *T* to complete the division. We know that in continuous illumination cells have a generation time of 6.5 h. The cells born in the long-day condition just after sunset (between 20:00 and 21:00) will completely mature in *N_1_* and arrest at the transition point during night.

Since *N_1_* is much shorter than the night duration (10 h) and total generation time of cells in light is ∼ 6.5 h (Fig. 2a) these cells are assumed to all be fully matured in *N_1_* and arrested at the transition point by sunrise at 6:00 the next morning. At 6:00, these cells will start to mature in *K_1_* (0.5 h), cross the transition point, mature in *N_2_* and divide. These cells will therefore divide *K_1_ + N_2_* after sunrise. Given that we can estimate *K_1_* (0.5 h) and measure the time at which the arrested cells divide the following morning (Extended Data Fig. 6b, 10:06), we can calculate *N_2_* as *N_2_* = time of division next morning - time of sunrise - *K_1_* ∼ 10:00 – 6:00 – 0.5 h ∼ 3.5 h *N_1_*: *N_1_* represents the time required for light-independent processes before a cell can cross the transition point. Since we know the total generation time of non-arrested cells (continuous illumination), *K_1_* and *N_2_* we can calculate *N_1_* = total generation time - *N_2_* = 6.5 h - 3.5 h = 3 h.

*σ*: The probability of a cell dividing at maturity *n* is represented as a Gaussian distribution centred around max(*K_1_*, *N_1_*) + *N_2_*, with a given width *σ*. The width of this distribution determines the variability of generation times experimentally observed. We estimated *σ* as 2 times the standard deviation around the mean generation time of cells growing in continuous illumination as *σ* = 2.86 h (Extended Data Fig. 6c). In the model, *σ* is discretized as 3 h.

*A_C_*: *A_C_* represents the chrysolaminarin accumulation rate during the light period. We estimate *A_c_* as the slope of a linear regression fitted to the chrysolaminarin data during a long day at 100 µmol m^-2^ s^-1^ = 5212 (chrysolaminarin fluorescence / h) (Extended Data Fig. 6d, Data same as in Fig. 4b).

*K_S_*: *K_S_* represents the Monod half velocity constant. We assume chrysolaminarin as a growth-limiting nutrient during darkness. In principle, cells can use both light and stored chrysolaminarin for energy production. Since cells born close to sunset, growing mostly in darkness, must depend mostly on chrysolaminarin for growth, we analysed the average growth rates of cells born at sunset or 2h before sunset across long-day and short-day conditions. We plotted the average growth rate (Data from single-cell growth experiments, Figs. 1-4) of these subset of cells against the corresponding levels of chrysolaminarin measured (Data from Fig. 4b). We fitted a Monod equation (Eq. (3)) and extracted the half velocity constant *K_s_* = 7102 (chrysolaminarin fluorescence) from these data. *K_S_* quantifies the chrysolaminarin concentration needed for a cell to grow at half the maximal growth rate achieved in continuous light exposure.

##### Conversion of chrysolaminarin accumulation rates to photosynthetic photon flux density

Our experimental data showed that the chrysolaminarin accumulation rates depend on light intensities. In order to test the impact of various light intensities on the daily growth rate of diatom populations (Fig. 6b), we inferred the chrysolaminarin accumulation rates at different light intensities. In a long day at 100 μmol m^−2^ s^−1^, the chrysolaminarin accumulation rate is 5212 a.u / h (Extended Data Fig. 6d). In a long-day at 43 μmol m^−2^ s^−1^, measuring maximal chrysolaminarin levels of 12’440 a.u. at the end of the light period (Fig. 4c) and assuming chrysolaminarin is fully depleted at the beginning of the light period in the long-day 43 μmol m^−2^ s^−1^ condition, we estimated an accumulation rate of 888 a.u. / h. At the light intensities investigated here (0-200 μmol m^−2^ s^−1^), the rate of photosynthesis in *T.pseudonana* is light limited and increases linearly with light intensities ^127^. Assuming the accumulation of chrysolaminarin is also light-limited between 0 and 200 μmol m^−2^ s^−1^, we established a linear relationship between light intensities and chrysolaminarin accumulation rates.

##### Calculation of cell ratios

In order to estimate the differences in biomass generated by populations growing with vs. without chrysolaminarin, we ran our simulations for 5 days and tracked the number of generated cells. At the end of the 5-day period, we calculated the ratio of the cumulative number of cells generated between populations growing with or without chrysolaminarin.

Due to the exponential nature of population growth, we estimated the number of cells after 10 days using population growth rate, rather than running our simulations for the entire 10 days. We first ran our simulations for a 5-day period. At the end of the 5-day period, we extracted the average generation time of cells within the population. From this, the number of cells at the end of a 10-day growth phase can be calculated as:

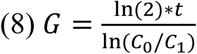

where *G* is the average generation time of a cell, *t* the duration of the growth phase and *C_0_* and *C_1_* the number of cells at the beginning and end of the growth phase respectively.

From the diatom bloom reports we cite in Fig. 6, we find that blooms can range from 6 to 20 days^76–78^. Furthermore, in a mesocosm bloom experiment, diatoms grew exponentially for 10 days^80^. Thus, a duration of 4 to 10 days seems a reasonable assumption for the duration of a diatom bloom. We further note that the scaling-up calculations can be easily adapted to other bloom durations.

##### Extraction of bloom parameters from satellite data

Six diatom blooms from the literature were selected, spanning different locations and seasons ^73–78^. From these reports, the coordinates and dates of the blooms were extracted. Using this information, the sunrise and sunset for a given location during a given time window can be calculated. From this, we then calculated the average diel light / dark cycles for each bloom.

Next, using data collected by the Moderate-Resolution Imaging Spectroradiometer (MODIS) of NASA’s Aqua satellite, we extracted the average Photosynthetically Active Radiation at the ocean surface ^128^ (PAR, µmol m^-2^ s^-1^) and the average diffuse attenuation coefficient for downwelling irradiance at 490 nm (*K_d_* 490, in m^-1^) ^129^ for each reported bloom. This was achieve by accessing the ocean color data (https://oceancolor.gsfc.nasa.gov/) acquired by the Moderate Resolution Imaging Spectroradiometer, accessed through GIOVANNI (https://oceancolor.gsfc.nasa.gov/resources/giovanni/). Further, we extracted the depth at which the bloom occurred, either directly from the bloom reports, or estimating it using the average DCM depth (Deep chlorophyll maxima) for the geographic location at which the bloom occurred^130^. Using Beer-Lambert law, we calculated the exponential extinction of PAR in the ocean for a given location and date with depth:

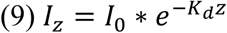

With *I_z_* the light intensity at depth *z*, *I_0_* the light intensity at the surface and *K_d_* the attenuation coefficient of PAR in the ocean water column for the given location and date reported for each bloom. The derived diel light / dark cycle and light intensities at depth for each bloom were then fed into our model to simulate growth of diatoms with and without chrysolaminarin storage under natural blooms conditions.

### Supplementary Videos

Supplementary Video 1

*T. pseudonana* cells growing in a 24 h light / 0 h dark cycle (Continuous day). Text top left indicates day / night cycle, timer top right indicates the duration of the experiment.

Supplementary Video 2

*T. pseudonana* cells growing in a 14 h light / 10 h dark cycle (Long day). Text top left indicates day / night cycle, timer top right indicates the duration of the experiment.

Supplementary Video 3

*T. pseudonana* cells growing in a 6 h light / 18 h dark cycle (Short day). Text top left indicates day / night cycle, timer top right indicates the duration of the experiment.

## Notes

### Competing Interest Statement

The authors have declared no competing interest.

